# The ALS/FTD-linked protein CHCHD10 associates with the TDP-43 C-terminal domain through a CHCH–helix interface

**DOI:** 10.64898/2026.08.01.742190

**Authors:** Karen S. Alarcón-Morales, Sara Pozo, Mauro Aguilera-Toste, Sara Martin-Ramos, Carolina Muguruza, Rodrigo Damian Garcia, Oscar Millet, Ganeko Bernardo-Seisdedos

## Abstract

Aggregation and cytoplasmic mislocalization of TDP-43 are defining features of several neurodegenerative disorders including Amyotrophic Lateral Sclerosis (ALS) and frontotemporal dementia (FTD). Yet the molecular interactions that regulate its transition from reversible assemblies to aggregation-prone states remain poorly understood. CHCHD10 is a mitochondrial protein genetically and pathologically linked to TDP-43 dysfunction, but the molecular basis connecting both proteins has remained unclear. Here, we combine solution Nuclear Magnetic Resonance (NMR) spectroscopy, biophysical assays and cellular imaging to characterize the interaction between human CHCHD10 and the C-terminal region of TDP-43 (TDP-43CTD). CHCHD10 comprises a dynamic N-terminal region and a folded CHCH domain that samples a reversible monomer–dimer equilibrium. Reciprocal NMR titrations show that the CHCH domain binds the conserved hydrophobic helix of TDP-43CTD through a dynamic submicromolar interaction that overlaps with self-association surfaces in both proteins. CHCHD10 alters the formation of ThT-reactive TDP-43CTD assemblies and reduces TDP-43CTD sedimentation under selected stoichiometric conditions, while itself becoming enriched in the sedimentable fraction. Equilibrium calculations, independently reproduced using a complete numerical mass-balance solution, identify the initial non-homodimeric CHCHD10 population as the strongest predictor of its subsequent sedimentation. In cells, full-length CHCHD10 shows stronger CHCH-dependent spatial association with TDP-43 than a construct lacking the CHCH domain. These findings define a CHCH–helix interface that couples homo- and heterotypic assembly equilibria and support an asymmetric interface-buffering model in which CHCHD10 can divert TDP-43CTD from self-association while increasing its own availability for recruitment into sedimentable assemblies.

## INTRODUCTION

TAR DNA-binding protein 43 (TDP-43) is an essential RNA-binding protein involved in RNA splicing, stability, transport and stress-granule dynamics. Under basal conditions, TDP-43 is predominantly nuclear, where its structured N-terminal domain, RNA-recognition motifs and low-complexity C-terminal region cooperate to support RNA metabolism and dynamic macromolecular assembly. In disease, however, TDP-43 undergoes nuclear depletion, cytoplasmic mislocalization and aggregation, defining a major pathological hallmark of ALS, FTD and a subset of Alzheimer’s disease cases (1). Although recent cryo-EM studies have resolved the architecture of mature TDP-43 amyloid filaments (2), the molecular interactions that regulate early TDP-43 assembly and prevent its conversion into pathological aggregates remain incompletely understood.

The low-complexity C-terminal region of TDP-43 (hereafter TDP-43CTD; residues 263–414) is central to this balance. This low-complexity region is intrinsically disordered but contains a highly conserved hydrophobic segment, commonly referred to as the conserved region, that transiently adopts α-helical structure and mediates helix-driven self-association. This region promotes TDP-43 condensation, nuclear retention and functional assembly, while also contributing to aggregation-prone behavior when the balance between reversible assembly and pathological conversion is disrupted (3, 4). Thus, the hydrophobic helix of TDP-43CTD represents both a functional interaction module and a vulnerable structural element whose misregulation may favor the emergence of TDP-43 proteinopathy.

Coiled-coil-helix-coiled-coil-helix domain containing 10 (CHCHD10, hereafter D10) is a small mitochondrial intermembrane-space protein genetically linked to ALS and FTD (5, 6). D10 has been implicated in mitochondrial cristae organization, synaptic integrity and stress responses (7), in part through its association with CHCHD2 and components of the mitochondrial contact site and cristae organizing system (MICOS) (8–10). Structurally, D10 comprises a dynamic N-terminal region followed by a conserved C-terminal CHCH domain containing the characteristic twin CX9C motif. The N-terminal segment displays low-complexity features and has recently been shown to form amyloid-like assemblies under defined conditions (11), whereas the CHCH domain is predicted to form a compact helical fold that may support protein–protein interactions and self-association.

Beyond its mitochondrial function, accumulating evidence links D10 to TDP-43 homeostasis (12). Depletion of D10 or expression of ALS/FTD-associated D10 variants promotes cytoplasmic TDP-43 accumulation, whereas wild-type D10 supports TDP-43 nuclear retention and protects against TDP-43-associated mitochondrial and synaptic defects (13–15). In human disease tissue, insoluble D10 aggregates have been detected alongside phosphorylated TDP-43 inclusions, and insoluble D10 levels correlate with insoluble TDP-43 burden (16). Recombinant studies further suggest that wild-type D10 can attenuate TDP-43 aggregate growth(14), while disease-associated D10 variants promote aggregation of both proteins (17). Together, these observations support a functional interplay between D10 and TDP-43, but whether this relationship is mediated by a direct molecular interaction, which domains are involved and how this interaction affects TDP-43 assembly remain unknown.

To address these questions, we used residue-resolved NMR together with complementary biophysical and cellular approaches to define the interaction between D10 and TDP-43CTD. We show that D10 comprises a flexible N-terminal region and a folded CHCH domain that samples a reversible monomer–dimer equilibrium. Reciprocal NMR titrations reveal that the CHCH domain directly engages the conserved hydrophobic helix of TDP-43CTD, coupling D10 self-association and TDP-43CTD recognition through overlapping interfaces. This interaction reshapes the assembly and sedimentation of both proteins in a concentration- and stoichiometry-dependent manner. Equilibrium analysis identifies the initial non-homodimeric D10 population as the strongest predictor of its subsequent recovery in sedimentable material, while heterotypic complex formation is associated with reduced TDP-43CTD sedimentation under selected conditions. In cells, full-length D10 shows stronger CHCH-dependent spatial association with TDP-43 than a construct lacking the CHCH domain. These findings establish a CHCH–helix interface that integrates D10 and TDP-43CTD within a common network of competing assembly equilibria.

## RESULTS

### D10 adopts a structured CHCH domain and a flexible N-terminus, sampling a monomer–dimer equilibrium

We first established a residue-level NMR framework to characterize the structural organization of human D10. Backbone resonances of full-length D10 were assigned to 92% completeness using a 150 μM sample recorded at 283 K under stabilizing reducing conditions containing 20 mM MES, pH 6.1, 150 mM NaCl, 5 mM DTT, 5 mM TCEP, 0.05% NaN_3_, 10% D_2_O and 2 M urea (BMRB 53350; Fig. 1A). Resonance identities were subsequently transferred through stepwise titrations to spectra recorded in the standard NMR buffer used for the subsequent experiments: 20 mM MES, pH 6.1, 150 mM NaCl, 0.05% NaN_3_ and 10% D_2_O at 308 K, without urea or reducing agents, unless otherwise indicated.

**Figure 1.**
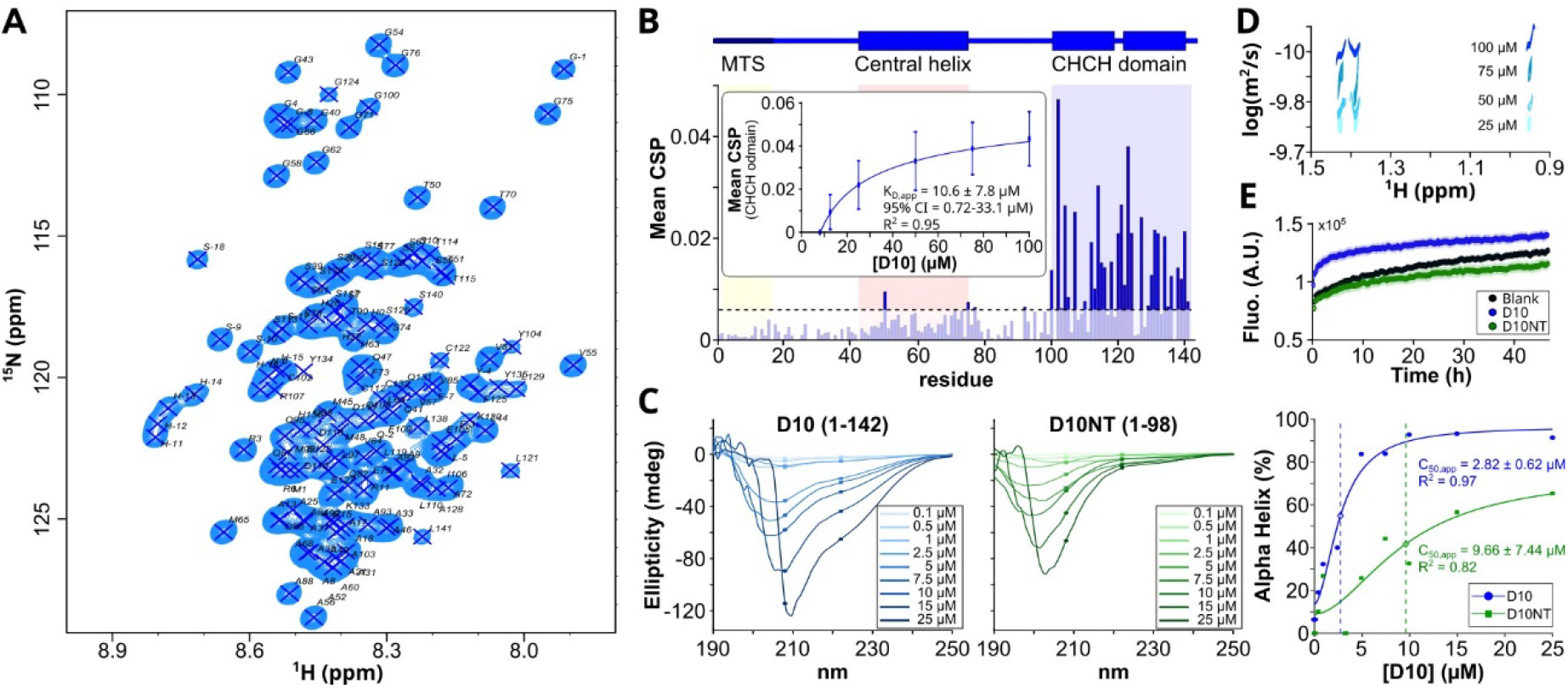
D10 contains a dynamic N-terminal region and a self-associating folded CHCH domain. **A**, Assigned ^1^H–^15^N HSQC spectrum of full-length human D10. Backbone resonances were assigned to 92% completeness using ^13^C/^15^N-labelled D10 recorded at 800 MHz and 283 K in 150 μM of protein, 20 mM MES pH 6.1, 150 mM NaCl, 5 mM DTT, 5 mM TCEP, 0.05% NaN_3_, 10% D_2_O and 2 M urea. Assigned residues are labelled. **B**, CSPs derived from concentration-dependent ^1^H–^15^N HSQC spectra of D10 recorded from 8 to 100 μM in 20 mM MES pH 6.1, 150 mM NaCl, 0.05% NaN_3_ and 10% D_2_O at 308 K. The lowest concentration spectrum was used as reference. The domain organization of D10 is shown above the plot, highlighting the mitochondrial targeting sequence (yellow), central helical region (red) and C-terminal CHCH domain (blue). The horizontal dashed line indicates the CSP threshold used to identify responsive residues. Inset, mean CSP as a function of D10 concentration. The global fit yielded K_D,app_ = 10.6 ± 7.8 μM, where the uncertainty corresponds to the standard error estimated from the local covariance matrix of the fit (R^2^=0.95); the profile-likelihood 95% confidence interval was 0.72–33.1 μM. Error bars represent the variability among the residues included in the global analysis. **C**, Far-UV circular dichroism spectra of full-length D10 and the isolated N-terminal construct D10NT recorded at increasing protein concentrations ranging from 0.1 to 25 μM. Proteins were prepared in 10 mM potassium phosphate pH 6.0 and spectra were acquired at 25 °C over the 190–250 nm wavelength range. Full-length D10 displayed a pronounced concentration-dependent α-helical signature, whereas D10NT retained a predominantly disordered spectral profile. Right, estimated α-helical content as a function of protein concentration. Empirical sigmoidal fitting yielded an apparent half-maximal concentration, C_50,app_ of 2.82±0.6 μM for D10 and 9.66±7.44 μM for D10NT. C_50,app_ denotes the protein concentration at which the fitted α-helical response reaches 50% of its maximum and should not be interpreted as a denaturation midpoint or thermodynamic dissociation constant. **D**, Diffusion-ordered NMR spectroscopy analysis of D10 at increasing concentrations. The progressive decrease in the apparent diffusion coefficient with increasing D10 concentration is consistent with an increase in average hydrodynamic size. **E**, Thioflavin-T fluorescence assay monitoring higher-order assembly of full-length D10 and D10NT. Proteins were incubated at 25 μM in 20 mM MES pH 6.1, 120 mM NaCl, 0.05% NaN_3_ and 15 μM ThT at 35 °C under quiescent conditions. Full-length D10 generated a stronger ThT-positive signal than D10NT or buffer control. Where shown, error bars or shaded regions represent variability between replicate measurements.

The ^1^H–^15^N HSQC spectrum of D10 revealed marked regional differences in chemical-shift dispersion, linewidth and signal intensity (Fig. 1A). This characteristic spectral partitioning was retained after transfer to the standard NMR buffer without urea. Resonances corresponding to the N-terminal region (residues 1–98) were intense and poorly dispersed, consistent with an intrinsically disordered segment. In contrast, residues belonging to the C-terminal CHCH domain (residues 99–142) displayed broader and more dispersed peaks with markedly reduced intensity, indicative of a folded domain experiencing slower molecular tumbling. Backbone secondary chemical shifts relative to random-coil values, particularly positive ΔδCα and ΔδCO values together with predominantly negative ΔδCβ and ΔδHα values, supported a continuous α-helical propensity across much of the CHCH domain (Supplementary Fig. S1) Circular dichroism measurements further supported this structural partitioning (Fig. 1C), where full-length D10 displayed a pronounced α-helical signature, whereas the isolated N-terminal region of D10 (hereafter D10NT; residues 1–98) exhibited a spectrum characteristic of predominantly disordered conformations. The concentration dependence of the estimated α-helical content, obtained using BeStSel (18), was described empirically using an apparent half-maximal concentration, C_50,app_, defined as the protein concentration at which the fitted α-helical response reached 50% of its maximum. This analysis yielded a C_50,app_ of 2.82 ± 0.62 μM for full-length D10 and 9.66 ± 7.44 μM for D10NT, although the latter estimate was comparatively poorly constrained. These apparent half-maximal concentrations provide an empirical description of the concentration-dependent CD response rather than a direct thermodynamic measure of folding or self-association.

Because disulfide-bond formation within the CHCH domain has been implicated in the mitochondrial import and stabilization of D10(19), we next assessed whether reducing conditions altered the structural properties of D10. ^1^H–^15^N HSQC spectra recorded in the presence of 5 mM DTT (incubated for > 2 h) were compared with spectra acquired under otherwise identical conditions without DTT (Supplementary Fig. S2A). The overall chemical-shift dispersion characteristic of the CHCH domain was retained under both conditions, arguing against widespread unfolding. Nevertheless, DTT-dependent chemical shift perturbations (CSPs) and intensity changes were concentrated predominantly within the CHCH domain, whereas the N-terminal region was comparatively weakly affected (Supplementary Fig. S2B,C). To determine whether these local NMR changes were accompanied by a loss of secondary structure, far-UV CD spectra were recorded for D10 and D10NT after incubation for more than 2 h in the presence of 1 mM TCEP and compared with matching samples prepared without TCEP (Supplementary Fig. S3A,B). The concentration-dependent α-helical signature of full-length D10 was largely preserved in the presence of TCEP, whereas D10NT retained a predominantly disordered spectral profile under both conditions. Together, these results indicate that reducing conditions perturb the local conformational or dynamic environment of the CHCH domain without causing detectable global loss of α-helical secondary structure.

We next asked whether the CHCH domain contributes to D10 self-association. Concentration-dependent ^1^H–^15^N HSQC spectra showed progressive and directional peak displacements as D10 concentration increased from 8 to 100 μM (Supplementary Fig. S4A). Representative residues within the CHCH domain, including C122, G124, Q131 and S140, followed continuous trajectories, consistent with fast exchange on the NMR timescale. Precisely, CSPs were strongly enriched in the CHCH region, whereas the N-terminal region displayed only minor changes (Fig. 1B and Supplementary Fig. S4B). The mean CSP increased with D10 concentration and approached saturation (Fig. 1B, inset), supporting a specific concentration-dependent equilibrium rather than nonspecific spectral drift.

To quantify this process, concentration-dependent CSPs from the 15 highest-ranked responsive residues were fitted globally using a mass-action monomer–dimer equilibrium model, with a single shared dissociation constant and residue-specific baselines and CSP amplitudes (see Materials and Methods). This analysis yielded an apparent dissociation constant of K_D,app_ = 10.6 ± 7.8 μM (fit estimate ± SE), and provided a good overall description of the concentration-dependent changes (R^2^ = 0.95; Fig. 1B, inset, and Supplementary Fig S5A). The asymmetric profile-likelihood 95% confidence interval was 0.72–33.1 µM, indicating that, although the central estimate was consistently in the low-micromolar range, its precise value remained comparatively weakly constrained.

The model predicted that the fraction of D10 protomers incorporated into dimers increased from approximately 0.45 at 8 μM to 0.79 at 100 μM, with the associated uncertainty shown in Supplementary Fig. S5C. The estimate was robust to the exclusion of individual resonances or concentration points: leave-one-residue-out and leave-one-concentration-out analyses produced median K_D,app_ values of 10.2 and 10.5 μM, respectively, with ranges of 5.3–24.5 and 6.7–22.4 µM. Residue-bootstrap analysis similarly yielded a median K_D,app_ of 12 μM, although the broad percentile distribution reflected the uncertainty expected for a weak and dynamic self-association process (Supplementary Fig. S5B).

To obtain an orthogonal readout of concentration-dependent self-association, Diffusion-Ordered SpectroscopY (DOSY) experiments were recorded at increasing protein concentrations. The apparent diffusion coefficient decreased progressively as the concentration increased from 25 to 100 μM, indicating an increase in the average hydrodynamic size of the protein population (Fig. 1D). Additional support for a dynamic self-association process was obtained from temperature-dependent NMR measurements. Upon increasing the temperature from 278 to 308 K, several CHCH-domain resonances sharpened and gained intensity, consistent with weakening of intermolecular contacts and a shift towards faster-tumbling species (Supplementary Fig. S6A,B). In contrast, resonances corresponding to the N-terminal region showed a modest decrease in intensity upon heating, likely reflecting enhanced conformational dynamics within this intrinsically disordered segment. Residue-specific temperature coefficients further highlighted the CHCH domain, together with parts of the central region, as the most temperature-sensitive elements of D10 (Supplementary Fig. S6B,C).

Finally, independently of the reversible equilibrium detected by NMR, we asked whether D10 could form ThT-positive higher-order assemblies during prolonged incubation. Full-length D10 generated a stronger ThT-positive signal than D10NT or the buffer control under quiescent conditions, whereas the isolated N-terminal fragment remained close to background throughout the assay (Fig. 1E). Because the NMR and ThT experiments probe markedly different timescales, these results do not establish that the reversible NMR-detected dimer is an obligate kinetic precursor of the ThT-positive assemblies. Rather, they show that the CHCH-containing full-length protein can access higher-order ThT-reactive states under the conditions tested, whereas the isolated N-terminal region is insufficient to reproduce this behavior.

### The CHCH domain of D10 engages the hydrophobic helix of TDP-43CTD through a dynamic submicromolar complex

To define the molecular interface between D10 and TDP-43, we performed reciprocal NMR titrations using either ^15^N-D10 or ^15^N-TDP-43CTD as reporters. Upon addition of unlabeled TDP-43CTD to ^15^N-D10, small but reproducible CSPs were detected almost exclusively within the folded CHCH region of D10, whereas the intrinsically disordered N-terminal segment remained largely unaffected (Fig. 2A,B and Supplementary Fig. S7A,B). The strongest perturbations clustered around residues L110, T114, C122, R127, A128, K130, Q131, C132 and Y134, defining a contiguous interaction surface that substantially overlaps with the self-association interface identified above.

**Figure 2.**
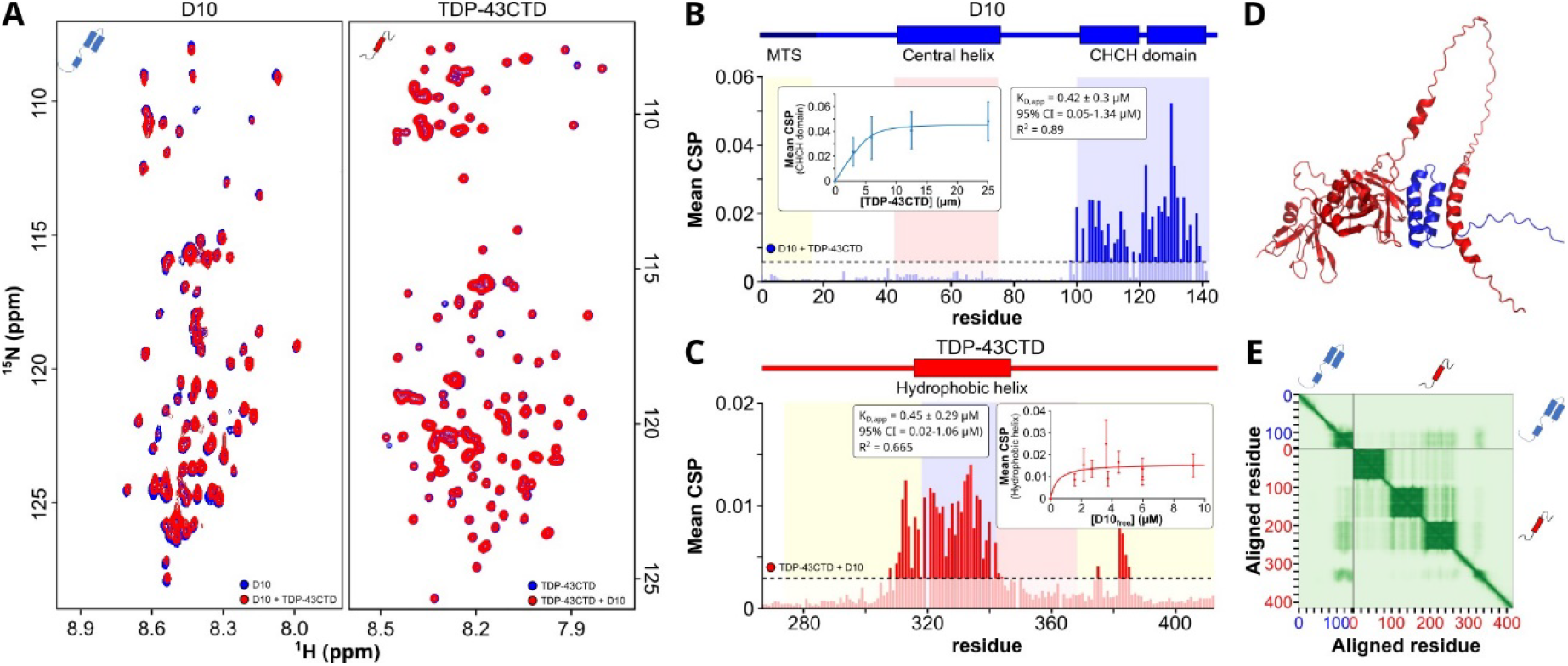
The CHCH domain of D10 engages the conserved hydrophobic helix of TDP-43CTD. **A**, Representative overlays of ^1^H–^15^N HSQC spectra from reciprocal NMR titrations between D10 and TDP-43CTD. Left, 12.5 μM of ^15^N-labelled D10 alone and after addition of unlabelled TDP-43CTD at 1:1 ratio. Right, 12.5 μM of ^15^N-labelled TDP-43CTD alone and after addition of unlabelled D10 at 1:1 ratio. Spectra were recorded in 20 mM MES pH 6.1, 150 mM NaCl, 0.05% NaN_3_ and 10% D_2_O at 308 K. **B**, CSPs in D10 upon addition of TDP-43CTD. Perturbations are strongly enriched in the C-terminal CHCH domain, whereas the N-terminal region and central helical segment remain only weakly affected. The domain organization of D10 is shown above the plot, with the mitochondrial targeting sequence, central helix and CHCH domain highlighted. Inset, mean D10 CSP as a function of TDP-43CTD concentration. CSPs were calculated from weighted ^1^H and ^15^N chemical shift differences. Dashed lines indicate the CSP threshold used to highlight perturbed residues. **C**, Reciprocal CSP mapping on ^15^N-labelled TDP-43CTD upon addition of D10. Perturbations cluster within the conserved hydrophobic helical region of TDP-43CTD and its immediate flanking residues. Inset, mean TDP-43CTD CSP as a function of the estimated monomeric D10 concentration, calculated from the D10 monomer–dimer equilibrium. **D**, AlphaFold3 model generated using full-length D10 and full-length TDP-43 without experimental restraints, templates or predefined interface information. The predicted complex positions the folded CHCH domain of D10 adjacent to the hydrophobic helical region of TDP-43CTD, in agreement with the reciprocal CSP-defined interaction surfaces. **E**, The PAE map is shown alongside the structural model. D10 is shown in blue and TDP-43 in red.

Because of this overlap, we considered whether the observed perturbations could simply reflect redistribution of D10 along its monomer–dimer equilibrium. However, for CHCH-domain resonances that report on both processes, the peak trajectories induced by addition of TDP-43CTD were clearly distinct from those observed upon increasing D10 concentration alone. This is evident for multiple shared reporters, including T114, C122, Q131, C132, Y134 and S140, which display different displacement directions and endpoints in the self-association and heterotypic titrations (compare Supplementary Figs. S4A and S7A). Thus, the TDP-43CTD-dependent CSPs do not merely reproduce the spectral signature associated with D10 homodimerization, although some contribution from redistribution among coupled homo- and heterotypic states cannot be excluded.

The D10 perturbations increased progressively with TDP-43CTD concentration and followed continuous peak trajectories, consistent with fast exchange on the NMR timescale (Supplementary Fig. S7A). The mean CSP approached saturation over the titration range (Fig. 2B, inset), supporting formation of a specific concentration-dependent complex. Fitting of the most responsive CHCH-domain residues yielded K_D,app_ = 420 ± 300 nM (estimate ± standard error; profile-likelihood 95% CI, 50–1340 nM; R^2^=0.892; RMSE = 0.006; Supplementary Fig. S8A). Parametric Monte Carlo propagation of the fitted residual variance yielded a median K_D,app_ of 420 nM and a 95% interval of 70–980 nM, providing an estimate of the uncertainty associated with residual noise under the fitted model (Supplementary Fig. S8B). Residue-bootstrap and leave-one-out analyses produced similar central estimates but broader distributions, consistent with the modest CSP amplitudes, limited titration range and dynamic nature of the interaction.

Reciprocal titrations monitored from the ^15^N-TDP-43CTD perspective revealed a complementary interaction pattern. Addition of D10 induced reproducible perturbations in TDP-43CTD without causing widespread spectral collapse (Fig. 2A). CSPs were concentrated primarily within the conserved hydrophobic helical region and its immediate flanking residues, with prominent effects across residues approximately 312–340, including F316, M322, Q327, A328, L330, S333, A334 and neighboring residues (Fig. 2C and Supplementary Fig. S9A,B). This region corresponds to the TDP-43CTD segment previously implicated in helix-mediated self-association, condensation and aggregation-prone behavior(3, 20, 21). In contrast, most residues outside this region showed only minor perturbations, although weaker changes were also detected around distal C-terminal residues, including the region surrounding A382-W385 (Supplementary Fig. S9A,B).

Across independent titration series recorded at different TDP-43CTD concentrations, the same hydrophobic-helix hotspot was consistently detected (Supplementary Fig. S9B). Global fitting of the CSP profiles from 12 responsive TDP-43CTD residues yielded K_D,app_ = 450 ± 290 nM (estimate ± standard error; profile-likelihood 95% CI, 20–1060 nM; R^2^=0.665; RMSE = 0.0048; Supplementary Fig. S10A), in close agreement with the value obtained from the D10-monitored titration. Parametric Monte Carlo propagation of residual uncertainty yielded a median K _D,app_ of 460 nM and a 95% interval of 1–1260 nM, reflecting the broader uncertainty of the TDP-43CTD-monitored dataset (Supplementary Fig. S10B). The central estimate remained stable upon residue-bootstrap and leave-one-residue-out analyses, whereas leave-one-condition-out analysis showed greater sensitivity to omission of individual titration conditions. Importantly, addition of an equimolar amount of the D10NT construct lacking the CHCH domain produced negligible CSPs and broadly preserved signal intensities across the sequence (Supplementary Fig. S11A–C). For both titration directions, the effective monomeric D10 concentration was estimated using the D10 homodimerization constant determined above. The reported K_D,app_ values and their confidence intervals are therefore conditional on a fixed D10 homodimerization K_D_ of 10.6 µM, and uncertainty in this parameter was not propagated into the binding fits. Consequently, the total uncertainty of the heterotypic interaction may be broader than the reported conditional intervals. Nevertheless, the close agreement between the independently monitored D10 and TDP-43CTD titrations provides internal consistency for an interaction centered in the submicromolar range.

Together, the reciprocal titrations independently defined complementary interaction surfaces comprising the folded CHCH domain of D10 and the conserved hydrophobic helical region of TDP-43CTD. To assess whether these experimentally mapped surfaces were structurally compatible within a common complex, we generated an AlphaFold3 model using full-length D10 and full-length TDP-43 without experimental restraints, templates or predefined interface information. As expected for a complex involving extensive intrinsically disordered regions, the global confidence metrics were modest (ipTM = 0.18; pTM = 0.23). Nevertheless, the highest-ranked model positioned the folded CHCH domain of D10 adjacent to the hydrophobic helical region of TDP-43CTD (Fig. 2D). The predicted interface overlapped closely with the two independently defined CSP hotspots: the CHCH-domain surface of D10 and the conserved hydrophobic helix of TDP-43CTD. Notably, this places TDP-43CTD on the same CHCH surface involved in D10 self-association, suggesting that TDP-43CTD recognition and D10 oligomerization are structurally coupled.

### D10 modulates TDP-43CTD assembly and is recruited into sedimentable material through its CHCH domain

Having established that the CHCH domain of D10 engages the aggregation-prone helical region of TDP-43CTD, we next examined how this interaction influences the assembly of both proteins in vitro. At 25 µM, TDP-43CTD alone showed a rapid increase in ThT fluorescence followed by a plateau (Fig. 3A). D10 altered this profile in a stoichiometry-dependent and non-monotonic manner: a substoichiometric amount of D10 increased the final ThT signal, whereas higher D10 concentrations progressively reduced it. The effects were less regular at 12.5 and 6.25 µM TDP-43CTD (Supplementary Fig. S12A,D,G). By contrast, D10NT produced only minor changes under equivalent conditions. Thus, full-length D10 modifies the formation or structural properties of ThT-reactive TDP-43CTD assemblies through its CHCH domain rather than acting as a simple concentration-dependent inhibitor.

**Figure 3.**
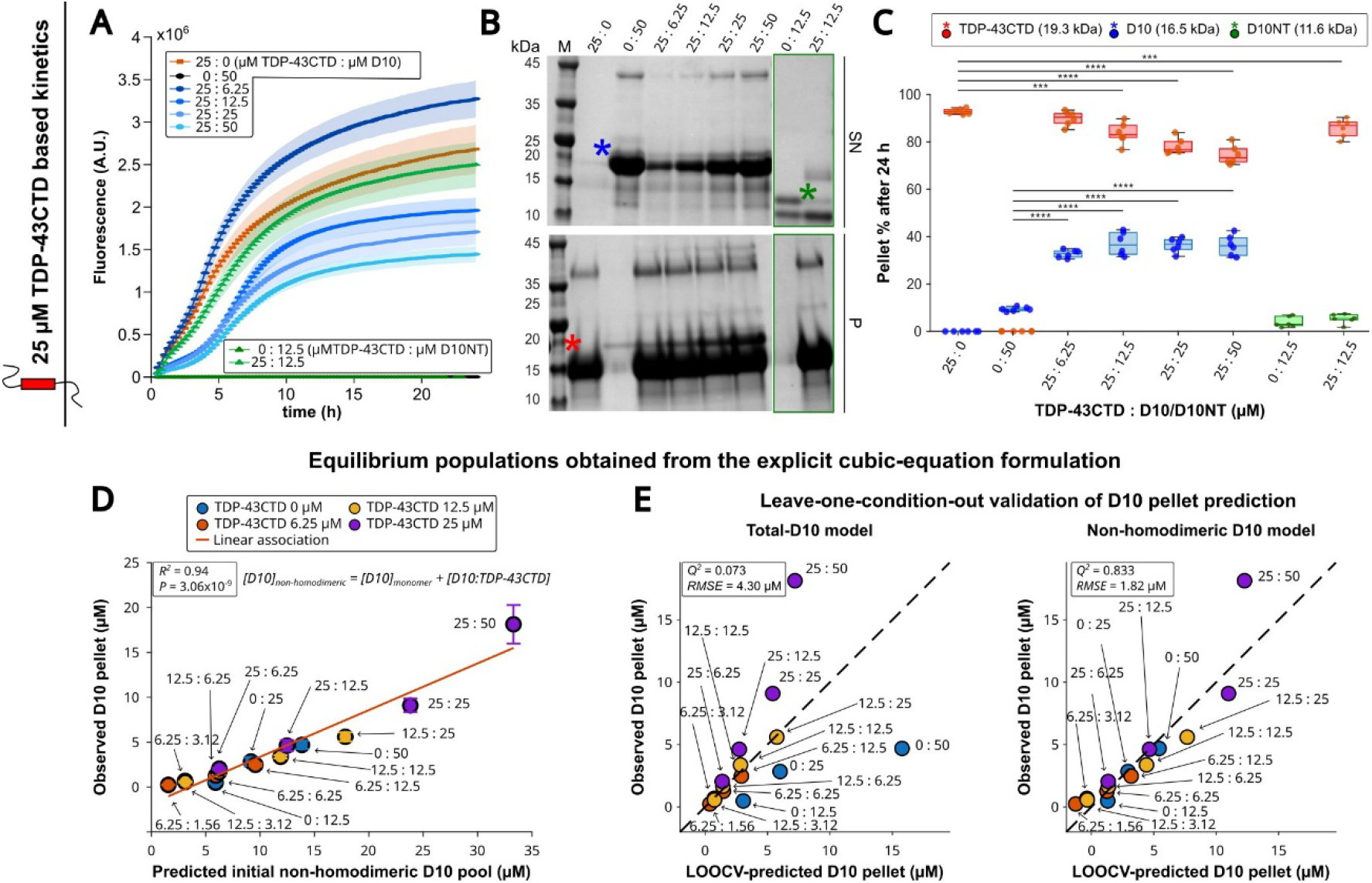
D10 reshapes TDP-43CTD assembly, and its initial non-homodimeric population predicts subsequent pellet recruitment. **A**, Thioflavin-T aggregation kinetics of 25 μM TDP-43CTD in the absence or presence of the indicated concentrations of full-length D10 or the isolated N-terminal construct D10NT. D10- and D10NT-only controls were included where indicated. Samples were prepared in 20 mM MES, pH 6.1, 120 mM NaCl, 0.05% NaN_3_ and 15 μM ThT, and incubated for 24 h at 35 °C under quiescent conditions. Full-length D10 produced concentration- and stoichiometry-dependent changes in TDP-43CTD ThT fluorescence, whereas D10NT had a comparatively minor effect. Lines represent mean fluorescence traces and shaded regions indicate variability between replicate measurements. **B**, Representative SDS–PAGE analysis of non-sedimented supernatant (SN) and pellet (P) fractions collected after 24 h. Samples were centrifuged at 16,000 × g for 30 min at 4 °C, and equivalent SN and P fractions were resolved on 4–12% polyacrylamide gels. Concentrations above the lanes are expressed as TDP-43CTD:D10 or TDP-43CTD:D10NT in μM. The D10NT-containing conditions are outlined in green. Colored asterisks indicate the bands quantified for TDP-43CTD (red; apparent molecular mass 19.3 kDa), D10 (blue; 16.5 kDa) and D10NT (green; 11.6 kDa). **C**, Densitometric quantification of the percentage of each protein recovered in the pellet after 24 h. TDP-43CTD was predominantly recovered in the pellet when incubated alone. Addition of D10 partially reduced TDP-43CTD pellet recovery under the selected conditions while markedly increasing D10 recruitment into the pellet. By contrast, D10NT remained predominantly non-sedimented and did not reproduce the reciprocal redistribution observed with full-length D10. Each point represents an independent replicate; boxes show the median and interquartile range. Statistical comparisons are indicated by brackets; NS, not significant; **P < 0.001 and ***P < 0.0001. **D**, Association between the calculated initial non-homodimeric D10 population and the experimentally observed amount of D10 recovered in the pellet after 24 h. Initial soluble populations were obtained from an explicit cubic-equation formulation incorporating D10 homodimerization (K_D,app_=10.6 µM) and formation of a 1:1 D10:TDP-43CTD complex (K_D,app_=0.434 µM), with TDP-43CTD treated as predominantly monomeric at the start of the reaction. The non-homodimeric D10 population was defined as [D10]_monomer_+[D10:TDP-43CTD]. Each point represents one initial TDP-43CTD:D10 condition and is colored according to the total TDP-43CTD concentration; labels indicate total TDP-43CTD:D10 concentrations in µM. Error bars indicate variability in the experimentally measured D10 pellet concentration. The red line represents the linear association over the experimentally sampled range (*R*^*2*^=0.938, *P*=3.06×10^−9^). **E**, Leave-one-condition-out cross-validation of linear models predicting D10 pellet recovery from either total D10 concentration or the calculated initial non-homodimeric D10 population. For each point, the indicated condition was excluded, the model was fitted to all remaining conditions and the excluded pellet value was predicted. Dashed lines indicate identity between predicted and observed values. The non-homodimeric D10 model showed substantially greater out-of-sample performance (*Q*^*2*^=0.833, *RMSE* = 1.82 µM) than the total-D10 model (*Q*^*2*^=0.073, *RMSE* = 4.30 µM). The equilibrium calculation describes the initial soluble populations and is not intended as a kinetic reconstruction of the 24-h sedimentation process.

Sedimentation analysis revealed a related but distinct effect. TDP-43CTD was predominantly recovered in the pellet when incubated alone. At 25 µM TDP-43CTD, pellet recovery decreased from 92.8 ± 1.1% in the absence of D10 to 83.5 ± 4.5%, 78.1 ± 3.3% and 74.4 ± 4.0% at TDP-43CTD : D10 ratios of 1:0.5, 1:1 and 1:2, respectively (Fig. 3B,C). In contrast, the 1:0.25 condition produced only a modest reduction in TDP-43CTD sedimentation despite yielding the highest ThT fluorescence. ThT fluorescence and pellet recovery therefore report different properties of the assemblies generated during the reaction. At lower TDP-43CTD concentrations, the effects of D10 on sedimentation were weaker and less monotonic, further indicating that the outcome depends on both protein concentration and stoichiometry (Supplementary Fig. S12F,I).

The redistribution was reciprocal. D10 alone was recovered only weakly in the pellet, whereas the presence of TDP-43CTD markedly increased D10 sedimentation (Fig. 3B,C). At 25 µM TDP-43CTD, D10 pellet recovery increased from 9.4 ± 0.9% in the D10-only condition to 32.7 ± 1.7%, 37.0 ± 4.9%, 36.4 ± 3.0% and 36.3 ± 4.3% across the four mixed conditions. Similar enrichment was observed in the 12.5 and 6.25 µM TDP-43CTD series. D10NT, however, remained predominantly non-sedimented both in the absence and presence of TDP-43CTD. The CHCH domain is therefore required not only for modulation of TDP-43CTD assembly but also for the recruitment of D10 into sedimentable material.

To relate this reciprocal redistribution to the soluble equilibria defined by NMR, we used the explicit cubic model described in Materials and Methods to calculate the initial populations of free D10 monomer, D10 homodimer and the 1:1 TDP-43CTD : D10 complex under each experimental condition. Total D10 concentration showed poor out-of-sample prediction of D10 pellet recovery, and neither free D10 monomer nor the heterotypic complex considered individually adequately described the complete dataset (Fig. 3E and Supplementary Fig. S13A). In contrast, the combined population of D10 outside the homodimeric state, defined as free D10 monomer plus D10:TDP-43CTD complex, showed a strong association with the amount of D10 recovered in the pellet after 24 h (R^2^ = 0.938, P = 3.06 × 10^−9^; Fig. 3D). This relationship retained substantial predictive performance under leave-one-condition-out validation (Q^2^ = 0.833, RMSE = 1.82 µM; Fig. 3E), remained robust to exclusion of complete TDP-43CTD concentration series and was preserved across the uncertainty ranges of the homo- and heterotypic dissociation constants (Supplementary Fig. S15A,B,D). An independent calculation using the complete coupled mass-balance system reproduced the same relationship (Supplementary Fig. S14).

The calculated populations showed that increasing TDP-43CTD redistributed D10 away from the homodimeric state and towards the heterotypic complex (Supplementary Fig. S15C). Consequently, conditions containing larger initial non-homodimeric D10 populations were also those in which more D10 was subsequently recovered in the pellet. Because the model describes the soluble populations at the beginning of the reaction, whereas sedimentation was measured after 24 h, this association does not identify a direct soluble precursor or reconstruct the temporal pathway of assembly. It nevertheless establishes that the initial partitioning of D10 between homodimeric and non-homodimeric states is informative about its subsequent sedimentation.

A complementary relationship was observed for TDP-43CTD. The calculated initial TDP-43CTD:D10 complex was associated with reduced TDP-43CTD pellet recovery relative to the corresponding D10-free controls (R^2^ = 0.806; Supplementary Fig. S13B). This relationship was driven predominantly by the 25 µM TDP-43CTD series and became weaker and non-monotonic at lower concentrations. Complex formation is therefore associated with reduced TDP-43CTD sedimentation under selected concentration regimes, rather than defining a general quantitative model of TDP-43CTD aggregation.

Together, these results reveal an asymmetric redistribution of the two proteins during assembly. D10 can reduce TDP-43CTD sedimentation under selected conditions, while TDP-43CTD expands the non-homodimeric D10 population that is most strongly associated with subsequent D10 recruitment into sedimentable material.

### The CHCH domain supports intracellular spatial association between D10 and TDP-43

Having established that the CHCH domain of D10 engages TDP-43CTD and reshapes its assembly *in vitro*, we next asked whether the presence of this domain influenced the intracellular spatial relationship between D10 and full-length TDP-43. HeLa cells were transiently transfected with TDP-43-Clover together with either full-length D10-Ruby or the N-terminal D10 fragment lacking the CHCH domain, D10NT-Ruby. Control conditions included Clover/Ruby, TDP-43-Clover/Ruby, Clover/D10-Ruby and Clover/D10NT-Ruby combinations. Representative fluorescence images showed partial co-distribution of full-length D10-Ruby and TDP-43-Clover across cytosolic and nuclear compartments, as well as within punctate condensate-associated regions (Fig. 4A).

**Figure 4.**
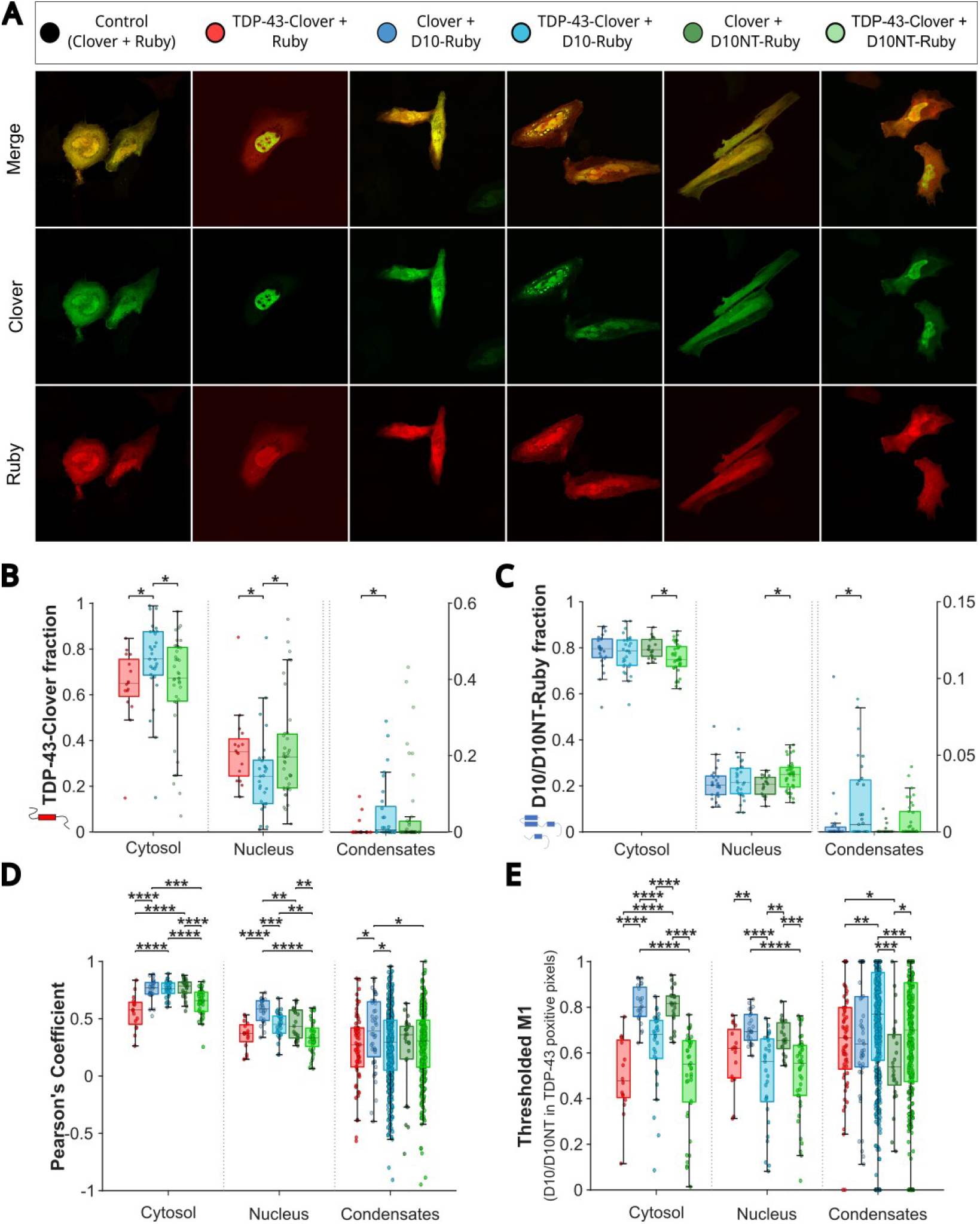
The CHCH domain supports intracellular spatial coupling between D10 and TDP-43. **A**, Representative confocal fluorescence images of HeLa cells transiently expressing the indicated Clover- and Ruby-tagged constructs. Control conditions included Clover/Ruby, TDP-43-Clover/Ruby, Clover/D10-Ruby and Clover/D10NT-Ruby. Interaction conditions included TDP-43-Clover co-expressed with either full-length D10-Ruby or the N-terminal construct D10NT-Ruby, which lacks the CHCH domain. Merge, Clover and Ruby channels are shown. Cells were fixed 18–24 h after transfection and imaged by confocal microscopy. **B**, Compartment-resolved distribution of TDP-43-Clover fluorescence across cytosolic, nuclear and condensate-associated regions. Co-expression with full-length D10-Ruby alters the cytosolic/nuclear distribution of TDP-43-Clover compared with TDP-43-Clover/Ruby, whereas D10NT-Ruby shows a weaker effect. **C**, Compartment-resolved distribution of D10-Ruby or D10NT-Ruby fluorescence across cytosolic, nuclear and condensate-associated regions. Condensate-associated fractions in B and C are plotted using the right y-axis. **D**, Pearson’s correlation coefficients calculated between Clover and Ruby signals in cytosolic, nuclear and condensate-associated regions. Full-length D10-Ruby shows higher spatial correlation with TDP-43-Clover than D10NT-Ruby in cytosolic and nuclear compartments. **E**, Thresholded Manders M1 analysis, reporting the fraction of D10-Ruby or D10NT-Ruby signal overlapping TDP-43-Clover-positive pixels. Full-length D10 shows increased overlap with TDP-43-positive regions compared with D10NT, particularly in cytosolic and condensate-associated compartments. Colocalization analyses were performed in Fiji/ImageJ using JaCoP after background subtraction. Regions of interest were defined for cytosolic, nuclear and condensate-associated compartments; low-quality ROIs were excluded before cell-level aggregation. Boxes indicate the interquartile range and median, and individual points represent cell-level measurements. Statistical comparisons were performed using Wilcoxon tests with FDR correction; *p < 0.05, **p < 0.01, ***p < 0.001, ****p < 0.0001.

We first analyzed the compartmental distribution of each fluorescent signal. Under the transient overexpression conditions used here, co-expression with full-length D10-Ruby shifted the distribution of TDP-43-Clover towards the cytosol. The median cytosolic fraction increased from 0.65 in cells expressing TDP-43-Clover/Ruby to 0.76 in cells expressing TDP-43-Clover/D10-Ruby, whereas the corresponding nuclear fraction decreased from 0.35 to 0.24 (False Discovery Rate (FDR)-adjusted p = 0.032 for both comparisons; Fig. 4B and Supplementary Fig. S17A). Co-expression with D10NT-Ruby produced a smaller change that did not reach statistical significance relative to TDP-43-Clover/Ruby. Moreover, the cytosolic and nuclear distributions of TDP-43-Clover differed significantly between the full-length D10 and D10NT co-expression conditions, further supporting a contribution of the CHCH-containing region to this phenotype (Fig. 4B and Supplementary Fig. S17A).

Conversely, TDP-43-Clover did not markedly alter the bulk cytosolic/nuclear distribution of full-length D10-Ruby. In contrast, D10NT-Ruby showed a modest shift towards the nuclear compartment upon TDP-43-Clover co-expression (FDR-adjusted p = 0.033; Fig. 4C and Supplementary Fig. S17B). Condensate-associated fractions were generally low but detectable. Full-length D10 co-expression increased the condensate-associated fraction of TDP-43-Clover relative to TDP-43-Clover/Ruby, whereas the corresponding effects on D10-Ruby and D10NT-Ruby distribution were comparatively modest (Fig. 4B,C and Supplementary Fig. S17A,B).

Because changes in bulk compartmental distribution do not establish a molecular interaction, we next quantified the spatial relationship between the Clover and Ruby signals using compartment-resolved Pearson’s correlation coefficients. In the cytosol, full-length D10-Ruby showed significantly higher correlation with TDP-43-Clover than D10NT-Ruby did (median Pearson’s coefficient, 0.764 versus 0.658; FDR-adjusted p = 2.52 × 10^−4^; Fig. 4D and Supplementary Fig. S18A). A similar CHCH-dependent difference was observed in the nucleus, where the full-length D10-Ruby/TDP-43-Clover combination showed higher correlation than D10NT-Ruby/TDP-43-Clover (median, 0.452 versus 0.333; FDR-adjusted p = 4.59 × 10^−3^). Within condensate-associated regions, Pearson’s coefficients were more heterogeneous and did not significantly distinguish full-length D10 from D10NT, consistent with the variability in size, intensity and composition of these punctate structures.

Directional Manders analysis further supported a CHCH-dependent difference in spatial overlap. Thresholded M1, reporting the fraction of D10-Ruby or D10NT-Ruby signal overlapping TDP-43-Clover-positive pixels, was higher for full-length D10 than for D10NT in the cytosol (median, 0.681 versus 0.551; FDR-adjusted p = 5.89 × 10^−3^) and in condensate-associated regions (median, 0.770 versus 0.699; FDR-adjusted p = 2.91 × 10^−3^; Fig. 4E and Supplementary Fig. S18C). Thus, a greater fraction of the full-length D10 signal occupied TDP-43-positive regions than was observed for the CHCH-deleted construct.

The reciprocal metric, thresholded M2, showed a more restricted pattern. The fraction of TDP-43-Clover signal overlapping D10/D10NT-positive pixels was significantly higher for full-length D10 than for D10NT in the nucleus, but not in cytosolic or condensate-associated regions (median nuclear M2, 0.698 versus 0.599; FDR-adjusted p = 3.72 × 10^−3^; Supplementary Fig. S18D). Consistent with this directional asymmetry, the symmetry index differed between full-length D10 and D10NT in nuclear and condensate-associated compartments (Supplementary Fig. S18B).

Together, these analyses show that the CHCH-containing full-length D10 protein exhibits a stronger intracellular spatial association with TDP-43 than the D10NT construct in this transient overexpression model. The clearest domain-dependent effects were observed in direct comparisons between full-length D10 and D10NT using Pearson’s and directional Manders coefficients. By contrast, the changes in bulk compartmental distribution represent related cellular phenotypes but cannot be attributed specifically to the NMR-defined interaction on the basis of these experiments alone. Although the imaging results are compatible with the CHCH-dependent molecular recognition and assembly behavior observed *in vitro*, they do not establish direct binding or a causal mechanism in cells.

## DISCUSSION

This study establishes a direct molecular link between D10 and an assembly-relevant region of TDP-43. The folded CHCH domain of D10 recognizes the conserved hydrophobic helix of TDP-43CTD, a segment that contributes to TDP-43 self-association, condensation and the transition towards higher-order assemblies. Importantly, this interaction does not occur between two otherwise independent proteins. The same CHCH surface participates in D10 homodimerization, whereas the TDP-43CTD helix is itself a self-association element. D10 and TDP-43CTD are therefore connected through a shared network of competing homo- and heterotypic interactions. This framework provides a mechanistic explanation for why D10 alters TDP-43CTD assembly without behaving as a conventional, uniformly acting aggregation inhibitor (Fig. 5).

**Figure 5.**
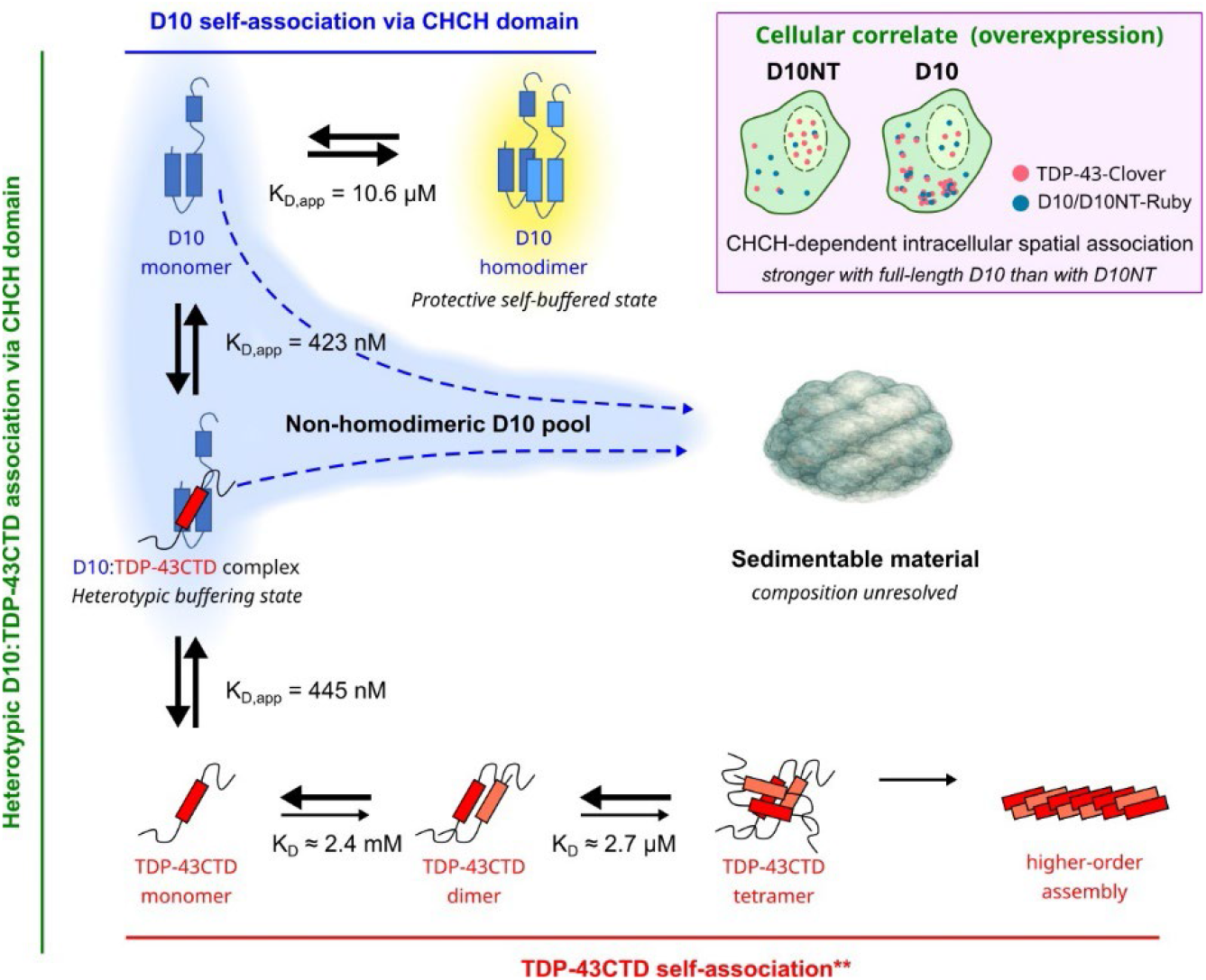
Working model of CHCH-dependent interface buffering and reciprocal redistribution of D10 and TDP-43CTD during assembly. D10 undergoes reversible self-association through its CHCH domain, with an apparent monomer–dimer dissociation constant of K_D,app_=10.6 μM. In the proposed model, occupation of the CHCH interface within the D10 homodimer provides a comparatively self-buffered state with reduced availability for alternative assembly interactions. Monomeric D10 also binds the conserved hydrophobic helix of TDP-43CTD to form a dynamic 1:1 heterotypic complex. The two values shown for this interaction (K_D,app_=423 and 445 nM) correspond to reciprocal NMR titrations monitored from the D10 and TDP-43CTD perspectives, respectively. Because the heterotypic interface overlaps with surfaces involved in the self-association of both proteins, complex formation is proposed to act as a transient interface-buffering state. **TDP-43CTD self-association is represented by the previously described sequential monomer–dimer–tetramer pathway, with an initially weak monomer–dimer transition (K_D_≈2.4 mM) followed by a more favorable dimer–tetramer transition (K_D_≈2.7 μM) considering previously reported values (4). Under the concentrations used in this study, TDP-43CTD is predicted to be predominantly monomeric at the beginning of the reaction. Binding of D10 to the TDP-43CTD helix may therefore reduce its access to the self-association pathway and divert part of the TDP-43CTD population away from sedimentable higher-order states. Conversely, heterotypic binding redistributes D10 outside the homodimeric population, expanding the non-homodimeric D10 pool, defined as free D10 plus D10:TDP-43CTD complex. This calculated population showed the strongest association with subsequent D10 recovery in the pellet. The dashed blue arrows indicate the association between the initial non-homodimeric D10 population and reciprocal redistribution into sedimentable material after 24 h. They do not specify the immediate assembly precursor or imply a resolved kinetic pathway. The composition, molecular architecture and route of formation of the final sedimentable assemblies remain unresolved. Yellow shading denotes putative buffering or relatively protected states, whereas the blue region denotes D10 populations outside the homodimeric reservoir. The inset summarizes the cellular correlate observed under transient overexpression: full-length D10 showed stronger CHCH-dependent spatial association with TDP-43 than the CHCH-deleted D10NT construct. The TDP-43CTD self-association constants were reported previously and were determined under experimental conditions different from those used here.

The mechanistic significance of the D10 interaction lies in this competition between interfaces. D10 samples a reversible monomer–dimer equilibrium through its CHCH domain, while TDP-43CTD engages an overlapping CHCH surface to form a dynamic heterotypic complex. The submicromolar apparent affinity obtained from reciprocal titrations, compared with the low-micromolar apparent affinity of D10 self-association, is consistent with TDP-43CTD efficiently engaging D10 molecules available outside the homodimer. These apparent constants should not be interpreted as a complete thermodynamic description of the system, but their relative magnitudes support a model in which D10 homodimerization and TDP-43CTD recognition compete within the experimentally sampled concentration range. The distinct NMR trajectories associated with the homo- and heterotypic interactions further indicate that TDP-43CTD does not simply shift the D10 monomer–dimer equilibrium, but forms a molecularly distinct complex through the same general interaction surface.

The equilibrium analysis suggests how this competition is translated into the reciprocal assembly behavior of the two proteins. D10 homodimerization can be viewed as a comparatively self-buffered reservoir in which occupation of the CHCH interface reduces its availability for alternative interactions. Consistent with this interpretation, the initial population of D10 outside the homodimeric state—comprising free D10 monomer and heterotypically bound D10—was more informative about subsequent D10 sedimentation than total D10 concentration or either soluble species considered separately. The relevant property is therefore not the abundance of a single obligate precursor, but the overall availability of D10 outside its homodimeric reservoir. This does not imply that the homodimer is inert or incapable of entering higher-order assemblies. Rather, homodimer occupancy appears to limit the population immediately accessible to competing assembly pathways.

The same heterotypic interaction has a different consequence for TDP-43CTD. The conserved helical region has recently been described as participating in a sequential self-association pathway in which the initial monomer–dimer transition is weak, whereas subsequent formation of higher-order helical assemblies becomes more favorable (4). Under the initial conditions examined here, TDP-43CTD is therefore expected to remain predominantly monomeric. Occupation of its hydrophobic helix by D10 may transiently restrict access to the first self-association step and divert part of this population away from sedimentable higher-order states. The association between heterotypic-complex formation and reduced TDP-43CTD sedimentation, most clearly observed at the highest TDP-43CTD concentration, is consistent with such an interface-buffering effect.

The overall outcome is thus asymmetric (Fig. 5). For TDP-43CTD, heterotypic binding can shield an assembly-driving surface and reduce access to sedimentable states. For D10, the same interaction competes with the comparatively protected homodimer and expands the non-homodimeric population associated with its later recruitment into the pellet. The heterotypic complex may therefore buffer the self-association interfaces of both proteins while occupied, yet produce opposite consequences at the level of their overall assembly landscapes. D10 diverts part of the TDP-43CTD population from sedimentable assembly, but does so at the cost of increasing its own availability for redistribution. This asymmetric cost of interface buffering reconciles the reduction in TDP-43CTD sedimentation with the simultaneous enrichment of D10 in sedimentable material.

This model also explains the non-monotonic ThT responses. If D10 acted simply by sequestering TDP-43CTD, increasing D10 concentration would be expected to produce a progressive reduction in assembly. Instead, the outcome depends on the relative populations of several competing states and therefore changes with concentration and stoichiometry. At substoichiometric D10 concentrations, partial occupation of the TDP-43CTD helix may alter nucleation, assembly morphology or accessibility of ThT-binding structures without substantially reducing the amount of sedimentable TDP-43CTD. At higher D10 concentrations, greater engagement of the helix may increasingly restrict entry into self-associated states. The divergence between ThT fluorescence and pellet recovery reinforces this interpretation: the two measurements report different properties of the assembling system and should not be treated as interchangeable measures of aggregation. Our data establish that D10 changes the assembly landscape of TDP-43CTD, but do not assign each fluorescence profile to a unique molecular species or pathway.

The inability of D10NT to reproduce these effects further identifies the CHCH domain as the organizing element of the interaction network. Although the low-complexity N-terminal regions of D10 and CHCHD2 can form amyloid-like assemblies under defined conditions (11), D10NT produced negligible NMR perturbations in TDP-43CTD, only modestly affected its ThT profiles and remained largely absent from the sedimentable fraction. The N-terminal region is therefore insufficient for TDP-43CTD recognition and reciprocal redistribution under the conditions examined here. The CHCH domain may additionally influence the assembly potential of the flexible N-terminus by controlling D10 self-association, partner recognition and local availability, even if the CHCH domain itself does not form the structural core of the final sedimentable material.

The proposed model provides a molecular framework for apparently contrasting observations linking D10 to TDP-43 homeostasis and disease. Wild-type D10 has been reported to attenuate TDP-43 aggregate growth, reduce phospho-TDP-43 pathology and rescue TDP-43-associated mitochondrial and synaptic defects (13, 14, 22, 23). At the same time, insoluble D10 has been detected alongside phosphorylated TDP-43 in human disease tissue, where the insoluble levels of both proteins are positively associated (16). Our results indicate that these observations need not be contradictory. D10 may transiently buffer an assembly-prone TDP-43 interface while itself becoming increasingly available for recruitment into insoluble material. Insoluble D10 associated with TDP-43 pathology could consequently represent failure or saturation of a protective interaction, pathological co-assembly, or the molecular cost of engaging aggregation-prone TDP-43 species.

This balance may be particularly sensitive to ALS/FTD-associated D10 variants. Mutations affecting CHCH-domain stability, D10 self-association or recognition of the TDP-43CTD helix could redistribute the equilibrium network in several ways. Destabilization of the D10 homodimer could expand the accessible non-homodimeric population, whereas weakened heterotypic recognition could reduce buffering of the TDP-43CTD helix. Conversely, an excessively stable or long-lived heterotypic interaction could promote persistent co-partitioning and reciprocal recruitment. Consistent with the latter possibility, D10^S59L^ has been reported to associate more strongly with TDP-43 and to promote TDP-43 insolubility and mitochondrial accumulation in cellular experiments (17). Disease-associated mutations may therefore alter the lifetime or consequences of the D10 interaction rather than simply eliminating it. The interface and equilibrium framework established here now provides a basis for distinguishing among these possibilities.

The cellular experiments provide a domain-dependent correlate of this molecular interaction. In the transient overexpression system, full-length D10 showed stronger spatial association with TDP-43 than D10NT, and a larger fraction of full-length D10 occupied TDP-43-positive regions (Fig. 5, inset). These observations support a contribution of the CHCH domain to intracellular recognition or co-partitioning, but do not demonstrate direct binding in cells or establish that the recombinant equilibrium model operates unchanged in the cellular environment. Full-length TDP-43, RNA, post-translational modifications, macromolecular crowding, compartmentalization and additional interaction partners will all influence the accessible states. Moreover, D10 participates in mitochondrial processes involving CHCHD2, MICOS organization, cristae maintenance and stress responses, providing several routes through which it could indirectly influence TDP-43 localization and proteostasis. Testing the CHCH–helix interaction in neuronal and disease-relevant models will therefore be necessary to determine how direct molecular recognition is integrated with the mitochondrial functions of D10.

The conserved TDP-43CTD helix is also emerging as a therapeutically actionable site. RNA chaperones that engage the RNA-recognition motifs of TDP-43 have been shown to allosterically remodel this region, promote aggregation-resistant conformations and reduce TDP-43 proteinopathy in cellular and animal models (24). The brain-penetrant small molecule XL20 similarly targets the conserved region, reduces TDP-43 neurotoxicity while largely preserving RNA-splicing activity and improves pathological outcomes in human ALS motor neurons and mouse models (25). Identification of the D10 CHCH domain as a natural binding partner for the same helix expands the range of molecular mechanisms capable of modulating this region. Structural definition of the CHCH–helix interface could eventually inform the development of peptides, miniproteins or small molecules that reproduce selected features of D10 recognition. However, the non-monotonic and asymmetric effects observed here also provide an important caution: productive target engagement will require control not only of binding affinity, but also of stoichiometry, interaction lifetime, subcellular localization and the assembly states generated after binding.

Several limitations define the scope of the proposed model. The recombinant experiments used TDP-43CTD rather than full-length TDP-43 and therefore exclude contributions from its N-terminal domain, RNA-recognition motifs and RNA binding. The apparent dissociation constants derive from dynamic, fast-exchanging systems and depend on the assumed equilibrium descriptions, while the published TDP-43CTD self-association constants were obtained under different experimental conditions and provide qualitative rather than directly comparable thermodynamic context. The AlphaFold3 model supports compatibility between the experimentally defined surfaces but does not establish the structure or dynamics of the complex. Most importantly, the equilibrium calculations describe the initial soluble populations, whereas sedimentation was measured after 24 h. They do not resolve the rates or sequence of the intervening transitions, determine whether the heterotypic complex remains soluble or dissociates before assembly, or establish whether D10 and TDP-43CTD occupy common sedimentable structures. Time-resolved measurements and direct structural analysis of the resulting assemblies will be required to answer these questions.

Together, our findings identify the CHCH domain of D10 and the conserved helix of TDP-43CTD as a shared regulatory interface connecting the self-association landscapes of both proteins. D10 homodimerization limits access to alternative interactions, whereas heterotypic recognition transiently buffers an assembly-driving surface of TDP-43CTD while expanding the D10 population available for redistribution. The resulting asymmetry explains why D10 can reduce TDP-43CTD sedimentation under selected conditions while itself becoming enriched in sedimentable material. More broadly, this work illustrates how competition between dynamic homo- and heterotypic interfaces can redirect aggregation-prone proteins without requiring irreversible sequestration. Whether this interface operates primarily as a protective buffer, a point of pathological vulnerability or both will depend on how its equilibrium is altered by cellular context and disease-associated mutations.

## MATERIALS AND METHODS

### Protein purification

Human D10 full-length (residues 1–142), the isolated N-terminal region of D10 lacking the CHCH domain (residues 1–98), and the C-terminal region of TDP-43 (residues 263–414), were used for recombinant protein experiments. Recombinant proteins were expressed as N-terminal His6-TEV fusion proteins in pET28A plasmid.

Recombinant proteins were expressed in Escherichia coli BL21(DE3). Bacterial cultures were grown in LB medium supplemented with kanamycin at 25 µg/ml at 37 °C with shaking until reaching an OD_600_ of 0.7–0.8. Protein expression was induced with 0.5 mM IPTG and continued overnight. Cells were harvested by centrifugation and bacterial pellets were stored at −80 °C until purification. Uniformly ^15^N-labelled and ^13^C/^15^N-labelled proteins were produced in M9 minimal medium following the same protocol.

Bacterial pellets were thawed on ice and resuspended in 50 ml of lysis buffer containing PBS 1×, 0.05% NaN_3_, 1 mM PMSF, cOmplete EDTA-free protease inhibitor cocktail and 10 µg/ml DNase I. Suspensions were incubated for 30 min at 4 °C with gentle agitation and lysed by sonication on ice using two 10-min cycles at 60% amplitude, with 20 s ON and 40 s OFF pulses. Lysates were centrifuged at 16000 g for 45 min at 4 °C, and soluble and insoluble fractions were collected separately.

Proteins were mainly purified from inclusion bodies. Inclusion body pellets were washed for 1 h at 4 °C in PBS 1× containing 1% Triton X-100, 500 mM NaCl, 10 µg/ml DNase I, 5 mM MgCl_2_ and 130 µM CaCl_2_, pH 7.8, followed by centrifugation at 16000 g for 45 min at 4 °C. A second wash was performed using the same buffer without DNase I, MgCl_2_ or CaCl_2_. The washed pellet was solubilized overnight at room temperature in denaturing buffer containing 20 mM Tris, 500 mM NaCl, 5 mM imidazole, 8 M urea and 0.05% NaN_3_, pH 7.8. Insoluble debris was removed by centrifugation and the clarified supernatant was filtered with a 0.45 μm filter before affinity purification.

Samples were loaded onto Ni-NTA agarose or HisTrap FF columns equilibrated in the appropriate buffer. Columns were washed with denaturing buffer and proteins were eluted with denaturing buffer supplemented with 500 mM imidazole. Eluted fractions were analyzed by SDS-PAGE, pooled and concentrated using Amicon Ultra centrifugal filters with a 3 kDa molecular weight cut-off. Protein purity was assessed on 4–12% precast polyacrylamide gels stained with BlueSafe Protein Stain.

TDP-43CTD was refolded by dialysis under acidic conditions (26). Denatured protein was dialyzed for overnight against 50 mM sodium acetate, pH 5.0, followed by a final 3 h dialysis step against 2.5 mM sodium acetate, pH 5.0. D10 and D10NT were refolded by dialysis or size-exclusion chromatography into buffers compatible with downstream experiments. For circular dichroism, proteins were prepared in 10 mM potassium phosphate, pH 6.0. For NMR and aggregation assays, proteins were prepared in MES-based buffers as described below. Although the constructs contained a TEV protease cleavage site, proteins were used in their His-tagged form because tag removal increased their aggregation propensity and have been proven not to be involved in the TDP-43 aggregation (21).

### Circular dichroism spectroscopy

Far-UV circular dichroism spectra were acquired to assess the secondary-structure content of D10 and D10NT. Proteins were prepared in 10 mM potassium phosphate, pH 6.0, and diluted to 0.1, 0.5, 1, 2.5, 5, 7.5, 10, 15 and 25 µM immediately before acquisition. Spectra were recorded at 25 °C using a Jasco J-815 spectropolarimeter equipped with a Peltier temperature-control system and continuous nitrogen flow. Measurements were performed in a 2 mm pathlength quartz cuvette over the 195–250 nm wavelength range, using a scan speed of 50 nm min−1, a bandwidth of 4 nm and a spectral resolution of 0.5 nm. Spectra were processed and plotted using MATLAB with the support of BeStSel (18).

### NMR spectroscopy

NMR spectra were acquired on a Bruker Avance III 800 MHz spectrometer equipped with a triple-resonance TCI cryoprobe. Unless otherwise indicated, ^15^N-labelled proteins were prepared in 20 mM MES, pH 6.1, 150 mM NaCl, 0.05% NaN3 and 10% D2O. Experiments were recorded at 308 K.

Backbone resonance assignments of D10 were obtained using ^13^C/^15^N-labelled D10 at 150 µM in 20 mM MES, pH 6.1, 150 mM NaCl, 5 mM DTT, 5 mM TCEP, 0.05% NaN3, 10% D2O and 2 M urea. Urea was included to reduce aggregation during the assignment experiments. A standard set of triple-resonance experiments was acquired, including HNCO, HNCA, HNCACB, HNcoCACB, HNHA and CON experiments, together with ^13^C-HSQC and ^15^N-HSQC spectra. Free induction decays were processed with NMRPipe(27) using apodization, phase correction and baseline correction. Peak picking, peak tracking and resonance assignment were performed using CcpNMR Analysis v3.3.2.3(28) and verified in TopSpin 3.2. DSS was used as the chemical shift reference for ^1^H, ^13^C and ^15^N. Backbone assignments were deposited in the BioMagResBank under accession code BMRB 53350.

Redox-dependent NMR experiments were performed using 100 µM D10. Non-reducing samples contained 20 mM MES, pH 6.1, 200 mM NaCl, 0.05% NaN3 and 10% D2O. Reduced samples were prepared in the same buffer supplemented with 5 mM DTT. Temperature-dependent ^1^H–^15^N HSQC experiments were recorded between 279 and 308 K using 50 µM D10.

### Analysis of D10 self-association by NMR chemical shift perturbations

To quantify the self-association behavior of D10 in solution, we analyzed the concentration dependence of backbone amide CSPs derived from ^1^H–^15^N HSQC spectra recorded at increasing protein concentrations (8, 12.5, 25, 50, 75 and 100 μM) under identical buffer conditions (20 mM MES pH 6.1, 150 mM NaCl, 0.05% NaN_3_, 10% D_2_O, 308 K).

Peak positions were extracted from manually curated assignments, and CSPs were calculated for each residue using the weighted chemical shift difference: 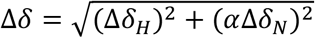, where Δδ_H_ and Δδ_N_ correspond to proton and nitrogen chemical shift differences, respectively, and α=0.2 accounts for the relative gyromagnetic scaling between nuclei. The lowest protein concentration (8 μM) was used as a reference spectrum.

To focus on residues reporting on the self-association process, the analysis was restricted to the CHCH domain (residues 99–142). Residues were filtered based on data quality criteria requiring (i) at least four valid CSP measurements across the titration series and (ii) a minimum CSP amplitude threshold. In addition, a monotonicity filter based on Spearman correlation between CSP and protein concentration was applied to retain residues displaying consistent concentration-dependent behavior.

To further assess the robustness of the analysis, residues were ranked according to their sensitivity to concentration-dependent changes, considering CSP amplitude, monotonicity, signal-to-noise proxy, and consistency across conditions. Global fitting was then repeated using subsets of the 15 top-ranked residues.

To quantify this process, CSP data were globally analyzed assuming a reversible monomer–dimer equilibrium 2*DD*10 ↔ *DD*10, with a dissociation constant defined as: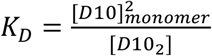, where and [D10_2_] represent the concentrations of monomer and dimer, respectively. Therefore, total protein concentration is given by *D*10_*tot*_= [*D*10]_*monomer*_+ 2[*D*10_2_]. Solving this system yields the monomer concentration, 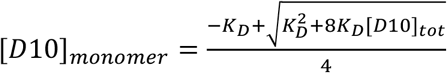, and the fraction of protomers in the dimeric state 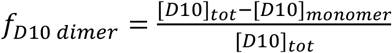 . The observed CSP for each residue_i_ at concentration [D10]_tot_ was globally modelled as Δ*δ*_*d*_ ([*D*10]_*tot*_ ) = *b*_*i*_ + [*f*_*D*10 *dimer*_ ([*D*10]_*tot*_ ; *K*_*D*_) − *f*_*D*10 *dimer*_ *C*_*ref*_ ; *K*_*D*_]Δ*δ*_*maxi*_ where: b_i_ is a residue-specific baseline offset accounting for systematic differences between spectra; Δδ_max,i_ is the residue-specific CSP amplitude; and C_ref_ corresponds to the reference concentration (8 μM).

All residues were fitted simultaneously using a global nonlinear least-squares approach, with a single shared dissociation constant K_D_ and residue-specific parameters b_i_ and Δδ_max,i_. The fitting procedure minimized the residuals between experimental and model-predicted CSP values across all residues and concentrations, excluding missing data points.

Optimization was performed using the Levenberg–Marquardt algorithm as implemented in MATLAB. The dissociation constant was parameterized in logarithmic space to ensure positivity and improve numerical stability. Initial values for residue-specific amplitudes were estimated from the maximum observed CSP for each residue, while baseline terms were initialized from median CSP values.

To evaluate the stability of the fitted parameters, resampling and subset analyses were performed. Bootstrap resampling of residues was used to estimate confidence intervals for K_D_, and additional fits were carried out after systematically excluding individual residues or concentrations. These analyses allowed assessment of the sensitivity of the fitted parameters to data selection and sampling variability.

Finally, fitted models were used to compute the fraction of protomers in the dimeric state as a function of total protein concentration. Residue-specific fits were inspected individually, and consistency across residues was assessed qualitatively and through comparison of normalized CSP profiles. Figures were generated using MATLAB, and all analyses were performed using custom scripts. The resulting dissociation constant is reported as an *apparent K*_*D*_, reflecting the underlying assumptions of a two-state monomer–dimer equilibrium and the experimental conditions employed.

### DOSY NMR

Diffusion-ordered NMR experiments were performed to assess concentration-dependent changes in the apparent translational diffusion coefficient of D10. Samples containing D10 at the concentrations indicated in Fig. 1D were prepared in the standard NMR buffer and analyzed at 308 K on an 800 MHz Bruker spectrometer. Experiments were acquired using the bipolar-gradient two-dimensional DOSY pulse sequence. Thirty-two gradient increments were collected using a sine-distributed gradient ramp from 2% to 98% of the maximum gradient strength. The diffusion delay (Δ) and gradient-pulse duration (δ) were set to 300 and 4 ms, respectively. Each gradient increment was acquired with 64 scans, 128 dummy scans and a recycle delay of 3 s. Data were processed in TopSpin using Fourier transformation, manual phase correction and baseline correction before DOSY reconstruction. Signal attenuation as a function of gradient strength was fitted to a single-component exponential decay using the standard dosy2d processing routine. Apparent diffusion coefficients were obtained from integrated D10 signals and are reported as population-averaged values. The measurements were used to compare relative changes in the hydrodynamic behavior of D10 across the concentration series and were not converted into absolute molecular masses.

### Reciprocal NMR titrations of D10 and TDP-43CTD

The D10/TDP-43CTD interaction was mapped using reciprocal ^1^H–^15^N HSQC titrations. In one set of experiments, ^15^N-labelled D10 was monitored upon addition of unlabeled TDP-43CTD. In the reciprocal experiment, ^15^N-labelled TDP-43CTD was titrated with unlabeled D10. Spectra were recorded in 20 mM MES, pH 6.1, 150 mM NaCl, 0.05% NaN3 and 10% D2O at 308 K.

CSPs and peak intensity changes were calculated from assigned ^1^H–^15^N HSQC spectra. Perturbed residues were mapped onto the corresponding protein sequence to identify the interaction surfaces. For D10-observed experiments, analysis focused on the CHCH domain, whereas for TDP-43CTD-observed experiments, perturbations were analyzed across the C-terminal TDP-43 fragment. Binding curves were fitted using a one-site binding model. Because D10 self-associates under the experimental conditions, effective monomeric D10 concentrations were estimated from the D10 monomer–dimer equilibrium and used for binding analysis. Apparent affinities are reported for the experimental conditions used.

### AlphaFold3 modelling

Structural modelling of the D10/TDP-43 interaction was performed using AlphaFold3. Full-length D10 and full-length TDP-43 sequences were provided as input without experimental restraints, templates or predefined interface information. Model confidence was evaluated using the default AlphaFold3 metrics, including pLDDT, PAE, ipTM and pTM. The predicted relative positioning of the D10 CHCH domain and the TDP-43 C-terminal hydrophobic helix was compared with the interaction surfaces identified by reciprocal NMR CSP analysis.

### Thioflavin-T aggregation assays and soluble–insoluble fractionation

Protein assembly was monitored by Thioflavin-T (ThT) fluorescence. Reactions were prepared in 20 mM MES, pH 6.1, 120 mM NaCl and 0.05% NaN_3_. The assembly behavior of D10 and D10NT was first examined independently at 25 µM. TDP-43CTD was analyzed at total concentrations of 6.25, 12.5 or 25 µM, either alone or in the presence of D10 or D10NT at the molar ratios indicated in the corresponding figures. Each reaction contained 15 µM ThT in a final volume of 50 µl and was prepared in a MicroAmp Optical 96-well reaction plate.

Fluorescence was recorded using a QuantStudio 5 Real-Time PCR System at 15-min intervals for 24 h. Reactions were maintained at 35 °C under quiescent conditions throughout the acquisition. Four wells were recorded for each experimental condition. For each well, the mean fluorescence of the first three acquisition cycles was used as the baseline and subtracted from the complete trace. Curves are presented as the mean fluorescence across the four wells, with variability represented by the corresponding standard deviation. Data processing and graphical representation were performed using custom MATLAB scripts. The ThT traces were analyzed as experimental readouts of the formation of ThT-reactive species and were not fitted to the equilibrium-population models described below.

At the end of the 24-h incubation, the four wells corresponding to each condition were pooled before fractionation. Samples were centrifuged at 16000 × g for 30 min at 4 °C to separate the non-sedimented supernatant and pellet fractions. Supernatants were collected, and pellets were washed once with assay buffer, re-centrifuged under the same conditions and resuspended in a volume of buffer equivalent to that of the corresponding supernatant. Samples were mixed with NuPAGE LDS Sample Buffer, heated at 95 °C for 10 min and resolved by SDS– PAGE using 4–12% precast polyacrylamide gels. Gels were stained with BlueSafe Protein Stain and imaged using an iBright CL1500 Imaging System.

Integrated band intensities were measured separately for TDP-43CTD and D10 or D10NT using ImageJ (29), allowing the distribution of each protein between soluble and insoluble fractions to be quantified after aggregation.

For each protein, the fraction recovered in the pellet was calculated as

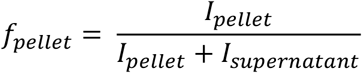

where I_pellet_ and I_supernatant_ are the corresponding densitometric signals. Pellet recovery was expressed either as a percentage, 100×f_pellet_, or as an absolute protein concentration, P_pellet_ = f_pellet_P_tot_, where P_tot_ is the total concentration of the protein in the initial reaction. Percentage values were used to compare protein partitioning experimentally, whereas absolute concentrations were used for the equilibrium-informed endpoint analyses.

Within each TDP-43CTD concentration series, pellet percentages were analyzed separately for TDP-43CTD and D10. Overall differences among conditions were assessed by one-way analysis of variance, followed by Tukey–Kramer multiple-comparison tests. For TDP-43CTD, the prespecified reference was the corresponding TDP-43CTD-only condition. For D10, mixed conditions were compared with the D10-only control included in the same concentration series. All reported P values were two-sided.

### Calculation of initial equilibrium populations using the explicit cubic formulation

The initial soluble populations present in each D10–TDP-43CTD reaction were calculated from the apparent equilibrium constants determined by NMR. The reduced model included D10 homodimerization, 2*DD*10 ⇌ *DD*10_2_, with the dissociation constant defined as 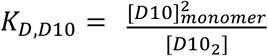, and formation of a 1:1 heterotypic complex between D10_monomer_ and monomeric TDP-43CTD, *D*10_*monomer*_ + TDP-43_CTD_ ⇌ D10:TDP-43CTD, with the dissociation constant defined as 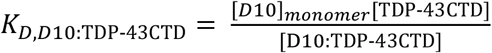.

Here, [D10]_monomer_, [D10_2_], [TDP-43CTD], and [D10:TDP-43CTD] denote the concentrations of free D10 monomer, D10 homodimer, free TDP-43CTD monomer, and the 1:1 heterotypic complex, respectively. The apparent dissociation constants were fixed at K_D,D10_=10.6 μM and K_D,D10:TDP43-CTD_=0.434 μM.

The corresponding mass-balance equations were: *D*10_*tot*_ = [*D*10]_*monomer*_ + 2[*D*10_2_] + [D10:TDP-43CTD] and TDP-43CTD_*tot*_ = [TDP-43CTD] + [D10:TDP-43CTD].

Using 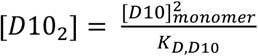 and 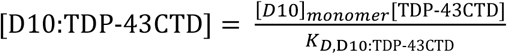, and eliminating the free TDP-43CTD monomer concentration through its mass-balance equation, the system was reduced to a single cubic equation in D10_monomer_:

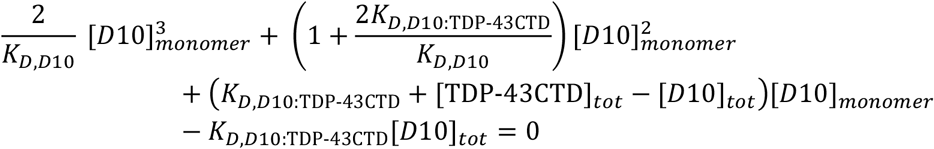

The three mathematical roots of the cubic equation were calculated numerically in MATLAB using the roots function. A root was considered physically admissible when it was real, non-negative, and no greater than [D10]_tot_. A single physically admissible root was obtained for every experimental condition.

Once [D10]_monomer_ had been determined, the remaining soluble populations were calculated as 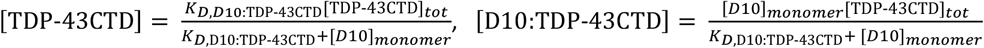, and 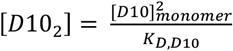.

Calculated solutions were accepted only when the absolute residuals of both protein mass-balance equations were below 10^−8^ μM. The concentration of D10 outside the homodimeric state was defined as: [D10]_non-homodimeric_ =[D10]_monomer_+[D10:TDP-43CTD].

Initial TDP-43CTD self-association was omitted from the reduced cubic formulation because the reported apparent monomer–dimer dissociation constant of approximately 2.4 mM is substantially greater than the 6.25–25 µM TDP-43CTD concentrations used here (Rizuan et al., 2025). The validity of this approximation was evaluated independently using the complete numerical mass-balance calculation described below.

### Numerical solution of the complete coupled mass-balance system

As an independent validation of the reduced cubic formulation, initial equilibrium populations were also calculated using a complete coupled model that retained D10 homodimerization, D10:TDP-43CTD complex formation and the reported sequential monomer–dimer–tetramer self-association equilibria of TDP-43CTD:

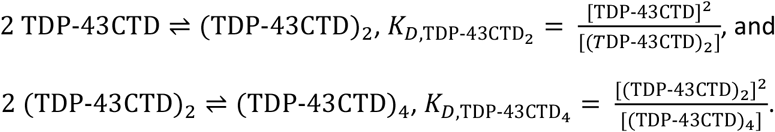

The fixed apparent dissociation constants used in this extended calculation were 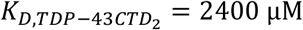 and 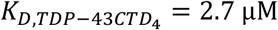, together with K_D,D10_=10.6 μM and K_D,D10:TDP-43CTD_=0.434 μM.

The complete D10 mass-balance equation was:

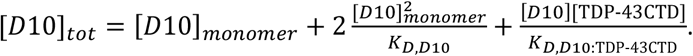

The complete TDP-43CTD mass-balance equation was:

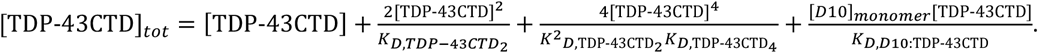

The two coupled nonlinear equations were solved for the free concentrations [D10]_monomer_ and [TDP-43CTD] using a custom damped-Newton algorithm implemented in MATLAB. Initial estimates were obtained from the analytical single-protein monomer–dimer solutions. At each iteration, the Newton correction was calculated from the analytical Jacobian of the two mass-balance residuals and applied using a damping factor of 0.7.

Free concentrations were constrained to the physical intervals 0≤[D10]≤[D10]_tot_ and 0≤[TDP-43CTD]≤ [TDP-43CTD]_tot_. Iteration was stopped when the maximum absolute mass-balance residual was below 10^−10^ μM, or after a maximum of 30 iterations.

The resulting free monomer concentrations were used to calculate the remaining equilibrium populations:

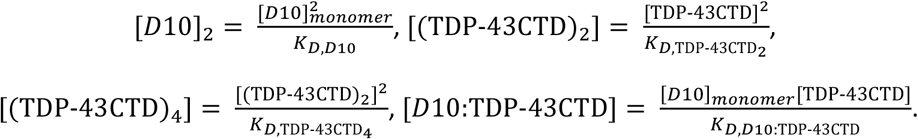

This extended calculation was used only to validate the initial equilibrium populations obtained from the explicit cubic formulation. It was not used to fit the ThT time courses, estimate aggregation-rate constants or reconstruct the temporal pathway leading to sedimentation.

### Equilibrium-informed analysis of the 24-h sedimentation endpoint

Associations between the calculated initial equilibrium populations and the amount of protein recovered in the pellet after 24 h were evaluated by ordinary least-squares linear regression with an intercept. For D10, the tested predictors were total D10 concentration, calculated free D10 monomer, predicted D10:TDP-43CTD complex and the non-homodimeric D10 pool, defined as [D10]+[ D10:TDP-43CTD]. The dependent variable was the experimentally measured absolute concentration of D10 recovered in the pellet.

Model performance in the complete dataset was summarized using the coefficient of determination, R^2^, and the root-mean-square error: 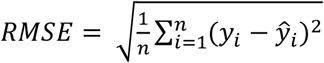.

The significance of each linear association was assessed using a two-sided Pearson correlation test.

Out-of-sample performance was evaluated by leave-one-condition-out cross-validation. Each experimental condition was excluded in turn, the regression was refitted using all remaining conditions and the pellet concentration of the excluded condition was predicted. Cross-validated performance was quantified using: 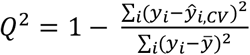 and the cross-validated RMSE. Robustness across concentration series was additionally assessed by leave-one-TDP-43CTD-series-out validation, in which all conditions containing 6.25, 12.5 or 25 µM TDP-43CTD were excluded together before refitting and prediction.

Sensitivity to uncertainty in the apparent equilibrium constants was assessed by recalculating all populations and repeating the regression and leave-one-condition-out analyses across the profile-likelihood confidence ranges of the D10 homodimerization and D10:TDP-43CTD dissociation constants.

For TDP-43CTD, the change in pellet recovery caused by D10 was calculated relative to the matched D10-free control: ΔT_pellet_ =T_pellet,control_ − T_pellet,+D10_.

Positive values therefore indicate a reduction in TDP-43CTD pellet recovery in the presence of D10, whereas negative values indicate increased recovery. The relationship between ΔT_pellet_ and the calculated initial D10:TDP-43CTD complex was analyzed by the same linear-regression and cross-validation procedures. Because this relationship differed among the three TDP-43CTD concentration series, it was interpreted as a concentration-dependent endpoint association rather than as a general kinetic model of TDP-43CTD aggregation. Conditions containing D10NT were used as experimental controls but were not included in the equilibrium calculations because no corresponding heterotypic binding equilibrium was detected by NMR.

### Cell culture and transient transfection

For cellular experiments, D10 and D10NT were expressed as Ruby fusion proteins, whereas TDP-43 was expressed as a Clover fusion protein. HeLa cells were maintained in DMEM high glucose without sodium pyruvate supplemented with 10% fetal bovine serum Advanced and 1% penicillin–streptomycin–fungizone. Cells were cultured at 37 °C in a humidified atmosphere containing 5% CO_2_ and used between passages 3 and 30. For transient transfection, approximately 150,000 cells were seeded per well in 6-well plates. After 24 h, cells were transfected with Lipofectamine 2000 using 2.5 µg of each plasmid per well. Cells were co-transfected with TDP-43-Clover together with either D10-Ruby or D10NT-Ruby. Eighteen to twenty-four hours after transfection, cells were fixed with 4% paraformaldehyde in PBS and stored in PBS at 4 °C. Samples were mounted using ProLong Glass Antifade Mountant, kept for 24 h at room temperature in the dark and then stored at 4 °C until imaging.

Fluorescence images were acquired using a Leica SP8 confocal microscope. 15 to 20 images were acquired per sample. Colocalization between Ruby- and Clover-labelled proteins was analyzed using the JACoP plugin in ImageJ (version 1.54p). Background was corrected before analysis using the Subtract Background function. Regions of interest were defined for whole-cell and subcellular analyses, including cytosolic, nuclear, nucleolar and condensate-associated regions. Pearson’s correlation coefficient and thresholded Manders overlap coefficients were calculated for each ROI. Thresholded M1 was defined as the fraction of D10-Ruby or D10NT-Ruby signal overlapping with TDP-43-Clover-positive pixels, whereas thresholded M2 was defined as the reciprocal fraction of TDP-43-Clover signal overlapping with D10-Ruby or D10NT-Ruby-positive pixels. ROIs with Quality scores below 9 were excluded. The remaining measurements were aggregated at the cell level before statistical analysis. Comparisons between D10-Ruby + TDP-43-Clover and D10NT-Ruby + TDP-43-Clover conditions were performed using Wilcoxon tests implemented in MATLAB.

In parallel, transfected cell samples were collected for protein extraction using RIPA. Lysis was performed in the same buffer. Supernatant and pellet fractions were separated. The pellet was resuspended in buffer containing 8 M urea. Protein samples were diluted in NuPAGE LDS Sample Buffer 4× and heated at 95 °C for 10 min. Proteins were resolved on 4–12% precast polyacrylamide gels using NuPAGE MOPS SDS running buffer and transferred to nitrocellulose membranes at 160 V for 1 h on ice using transfer buffer containing 20% ethanol. Membranes were washed three times with PBS containing 0.1% Tween-20, blocked for 1 h at room temperature with 5% Blotting-Grade Blocker in TPBS and incubated overnight at 4 °C with primary antibodies diluted in blocking buffer. Membranes were then washed three times with TPBS, blocked again for 10 min and incubated for 1 h at room temperature with the corresponding HRP-conjugated secondary antibodies. Bands were detected using SuperSignal West Pico PLUS chemiluminescent substrate and imaged with an iBright CL1500 imaging system.

The primary antibodies used were anti-D10 (Sigma, HPA003440), anti-TDP-43/TARDBP (Abnova, H00023435-M01), anti-His (Abyntek, ABK1-A24249) and anti-β-actin (Cell Signaling Technology, 8H10D10 #3700). HRP-conjugated goat anti-rabbit IgG (Jackson Immuno Research, 111-035-144) and rabbit anti-mouse IgG (Jackson Immuno Research, 315-035-045) were used as secondary antibodies.

## ACKNOWLEDGMENTS

The authors thankfully acknowledge the NMR resources and technical support provided by the Laboratorio de RMN de Euskadi (LRE) at CIC bioGUNE, a node of the Spanish ICTS Red de Laboratorios de RMN de Biomoléculas (R-LRB). The authors are particularly grateful to Dr. Tammo Diercks for his invaluable technical assistance with NMR data acquisition and for his expert advice throughout the project.

## DATA AVAILABILITY

Backbone NMR assignments for D10 have been deposited in the Biological Magnetic Resonance Data Bank under accession code BMRB 53350. Additional data and custom MATLAB scripts supporting the findings of this study are available from the corresponding author upon reasonable request.

