## Supplementary Material for "The ALS/FTD-linked protein CHCHD10 associates with the TDP-43 C-terminal domain through a CHCH–helix interface"

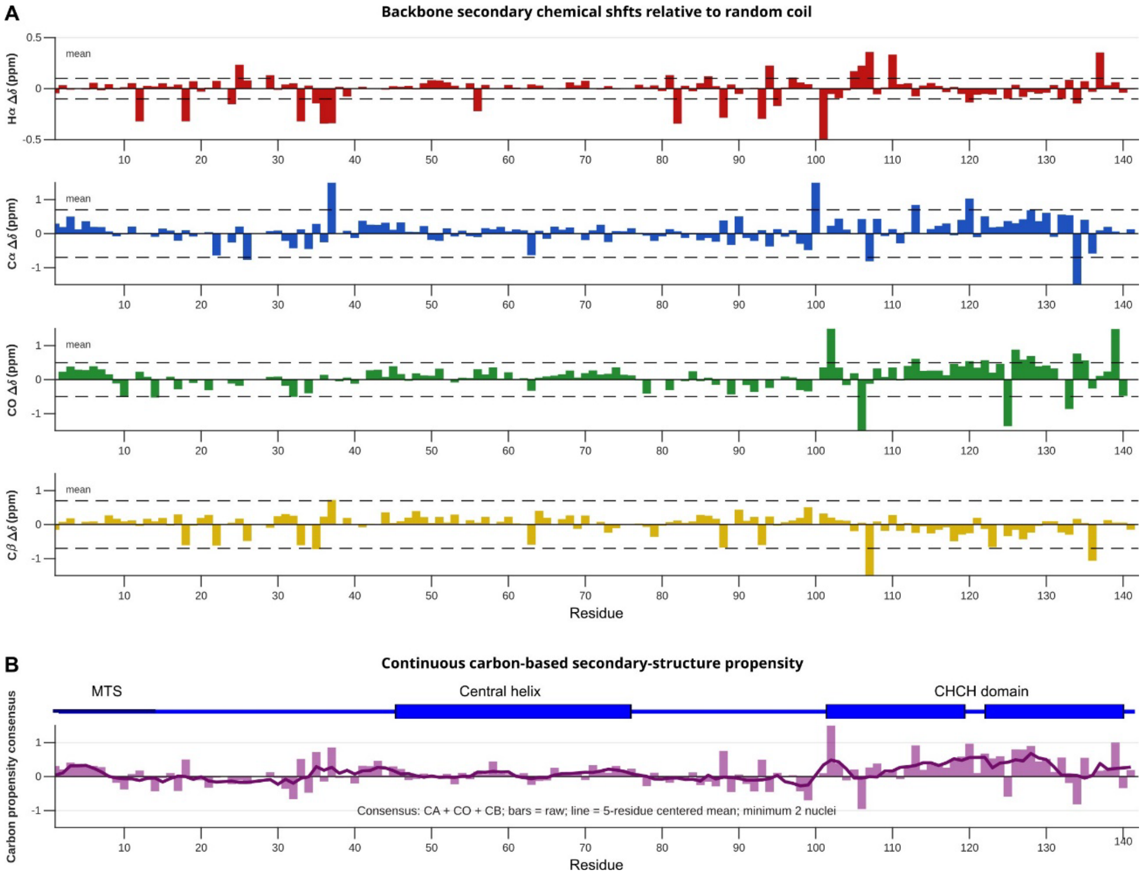


**Supplementary Fig. S1. Backbone secondary chemical shifts and residue-level secondary-structure propensity of D10. A,** Residue-specific secondary chemical shifts of Hα, Cα, CO and Cβ nuclei, calculated relative to random-coil reference values. Positive ΔδCα and ΔδCO values, together with predominantly negative ΔδCβ and ΔδHα values, are consistent with α-helical conformational propensity, particularly within the C-terminal CHCH domain. Horizontal dashed lines indicate the positive and negative mean absolute secondary chemical shift for each nucleus and are included as visual references. **B**, Continuous carbon-based secondary-structure propensity calculated from the combined Cα, CO and Cβ secondary chemical shifts, requiring data from at least two nuclei per residue. Bars represent the raw residue-level consensus values, whereas the solid line represents a five-residue centred moving average. Positive values indicate α-helical propensity, while negative values indicate extended or β-like conformational propensity.

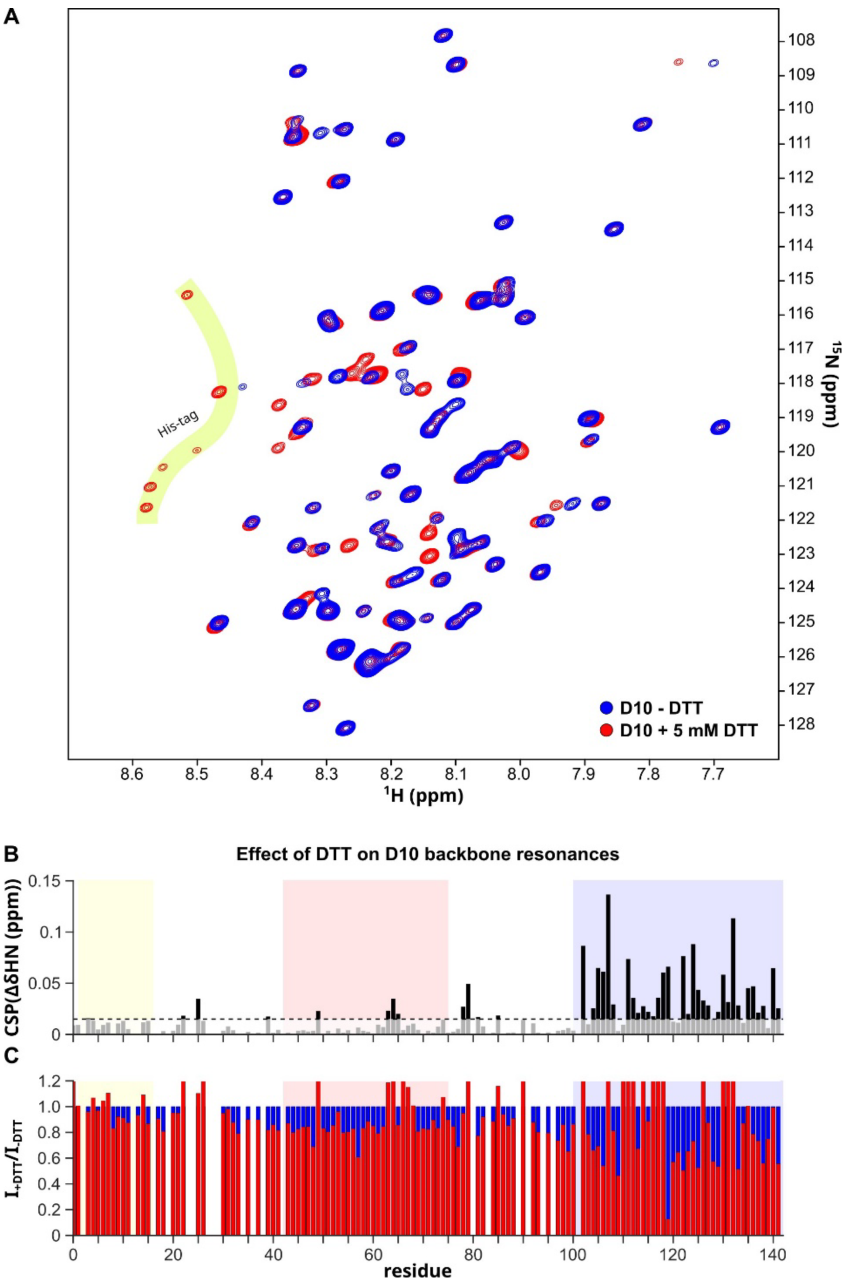


**Supplementary Figure S2. Effect of DTT on the NMR spectral properties of D10. A**, Overlay of $^{1}$H–$^{15}$N HSQC spectra of D10 recorded under otherwise identical conditions in the absence of DTT (blue) or in the presence of 5 mM DTT (red) incubated with the protein for > 2 h. The overall chemical-shift dispersion was retained under both conditions, although a subset of resonances displayed DTT-dependent chemical-shift and intensity changes. Resonances attributed to the N-terminal His-tag are indicated and were excluded from the residue-resolved analyses in panels B and C. **B,** Residue-specific backbone amide chemical shift perturbations between the DTT-free and DTT-containing conditions. Perturbations were concentrated primarily within the C-terminal CHCH domain, whereas the N-terminal region was comparatively weakly affected. **C,** Residue-specific peak-intensity ratios, calculated as $I_{+DTT}/I_{-DTT}$. The horizontal line at 1 indicates unchanged intensity between conditions; values above or below 1 indicate signal enhancement or attenuation, respectively, upon addition of DTT.

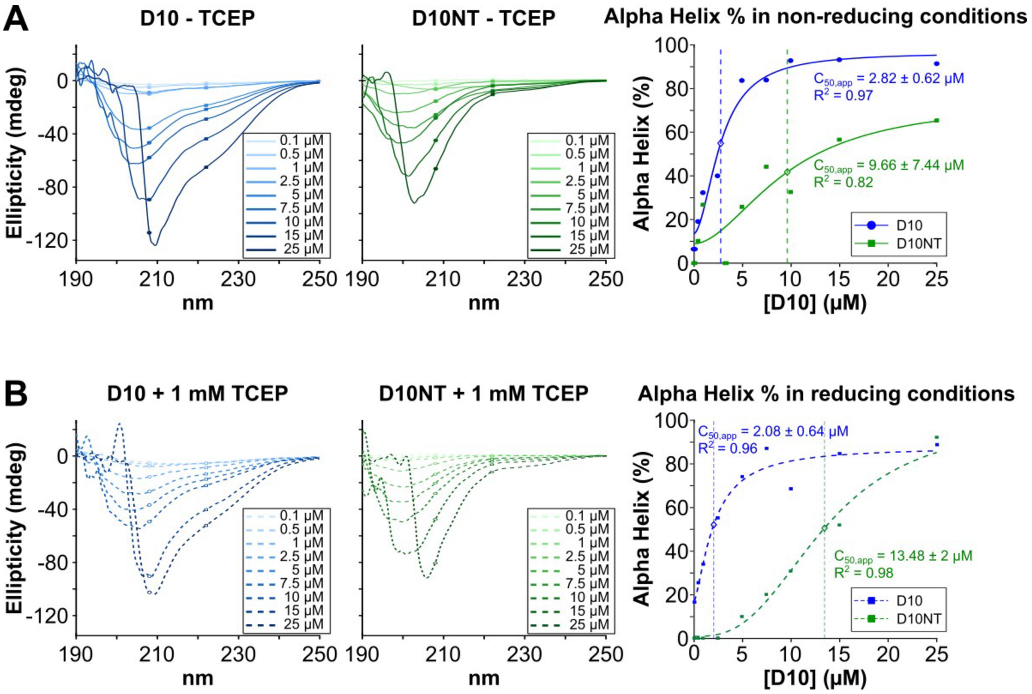


**Supplementary Fig. S3. Far-UV circular dichroism analysis of D10 and D10NT in the absence and presence of TCEP. A,** Far-UV CD spectra of full-length D10 and the isolated N-terminal construct D10NT, comprising residues 1–98, recorded at protein concentrations ranging from 0.1 to 25 µM in the absence of reducing agent. Right, estimated α-helical content as a function of protein concentration. Solid lines represent empirical sigmoidal fits, and $C_{50,app}$ denotes the protein concentration at which the fitted α-helical response reaches 50% of its maximum. **B,** Equivalent CD measurements performed after incubation of D10 and D10NT for more than 2 h in the presence of 1 mM TCEP. The concentration-dependent α-helical signature of full-length D10 was largely preserved in the presence of TCEP, whereas D10NT retained a predominantly disordered spectral profile under both conditions. Fit parameters are indicated in the corresponding plots. $C_{50,app}$ provides an empirical description of the concentration dependence of the CD-derived α-helical response and should not be interpreted as a denaturation midpoint or thermodynamic dissociation constant. Spectra were recorded in 10 mM potassium phosphate, pH 6.0, at 25 °C over the 190–250 nm wavelength range.

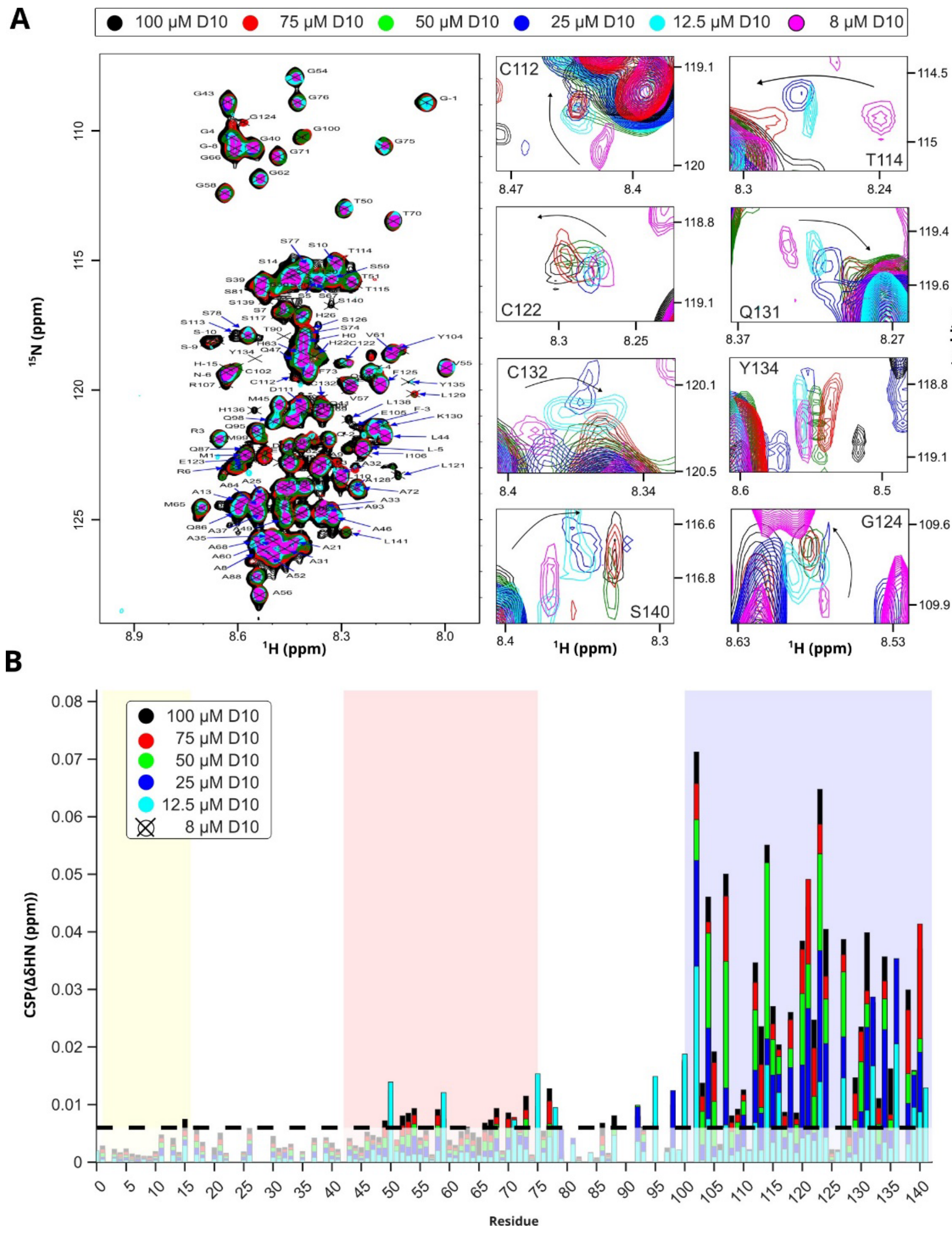


**Supplementary Fig. S4. Concentration-dependent NMR analysis of D10 self-association. A**, Overlay of $^{1}H$–$^{15}N$ HSQC spectra of full-length D10 recorded at increasing protein concentrations. Spectra were acquired at 8, 12.5, 25, 50, 75 and 100 µM $^{15}N$-labelled D10 in 20 mM MES pH 6.1, 150 mM NaCl, 0.05% $NaN_3$ and 10% $D_2O$ at 308 K. Representative CHCH-domain residues, including C122, G124, Q131 and S140, show continuous concentration-dependent peak trajectories, consistent with fast exchange on the NMR timescale. **B**, CSPs calculated relative to the lowest D10 concentration. Perturbations increase with D10 concentration and are strongly enriched in the CHCH domain, whereas the N-terminal region displays only minor changes. The shaded regions indicate the mitochondrial targeting sequence, central helical region and CHCH domain.

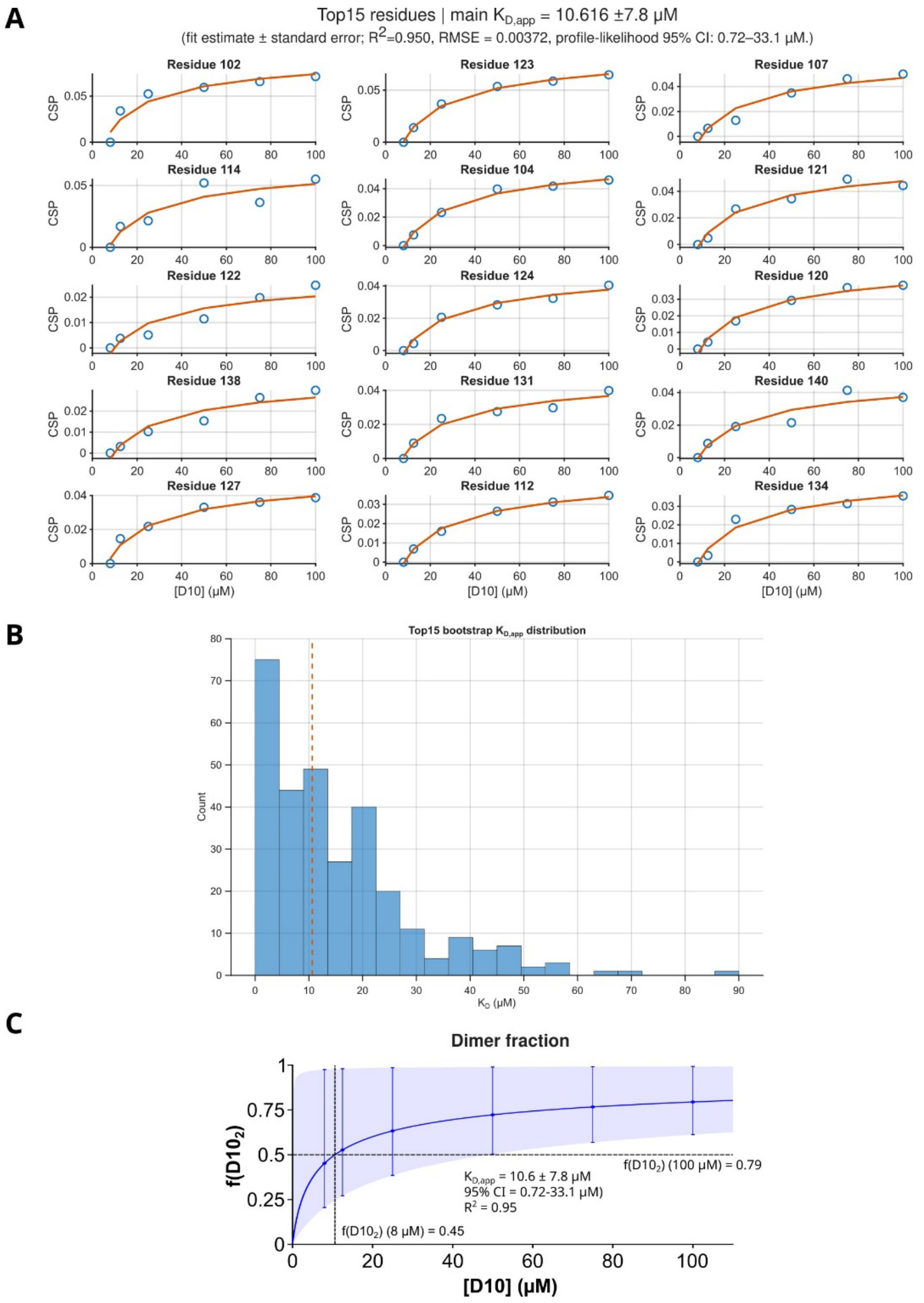


**Supplementary Fig. S5. Robustness of the monomer–dimer fitting analysis for D10 self-association. A**, Individual concentration-dependent CSP profiles for the 15 most responsive residues used to evaluate the robustness of the D10 self-association fit. Experimental CSP values are shown together with fits to a reversible monomer–dimer equilibrium model. Residues within the CHCH-containing region show coherent saturable behavior , supporting a shared concentration-dependent self-association process. **B**, Bootstrap distribution of apparent $K_D$ values obtained from resampling of the top-ranked residues. The dashed line indicates the global fitted $K_{D,app}$ = 10.6 ± 7.8 µM (fit estimate ± standard error; $R^2$=0.950, RMSE = 0.00372). The profile-likelihood 95% confidence interval was 0.72–33.1 µM. The broad distribution reflects the weak and dynamic nature of the equilibrium and the uncertainty inherent to CSP-based fitting. **C**, Predicted fraction of D10 protomers incorporated into homodimers, $f_D = 2[D10_2]/[D10]_{total}$, as a function of total D10 concentration, calculated using the fitted monomer–dimer equilibrium. The solid line represents the prediction obtained with the central KD,app estimate, whereas the shaded region and error bars show the uncertainty propagated from the profile-likelihood confidence interval. The model predicts an increase in the dimer-associated protomer fraction from approximately 0.45 at 8 µM D10 to 0.79 at 100 µM D10. The vertical dashed line marks the fitted $K_{D,app}$, and the horizontal dashed line indicates a dimer fraction of 0.5.

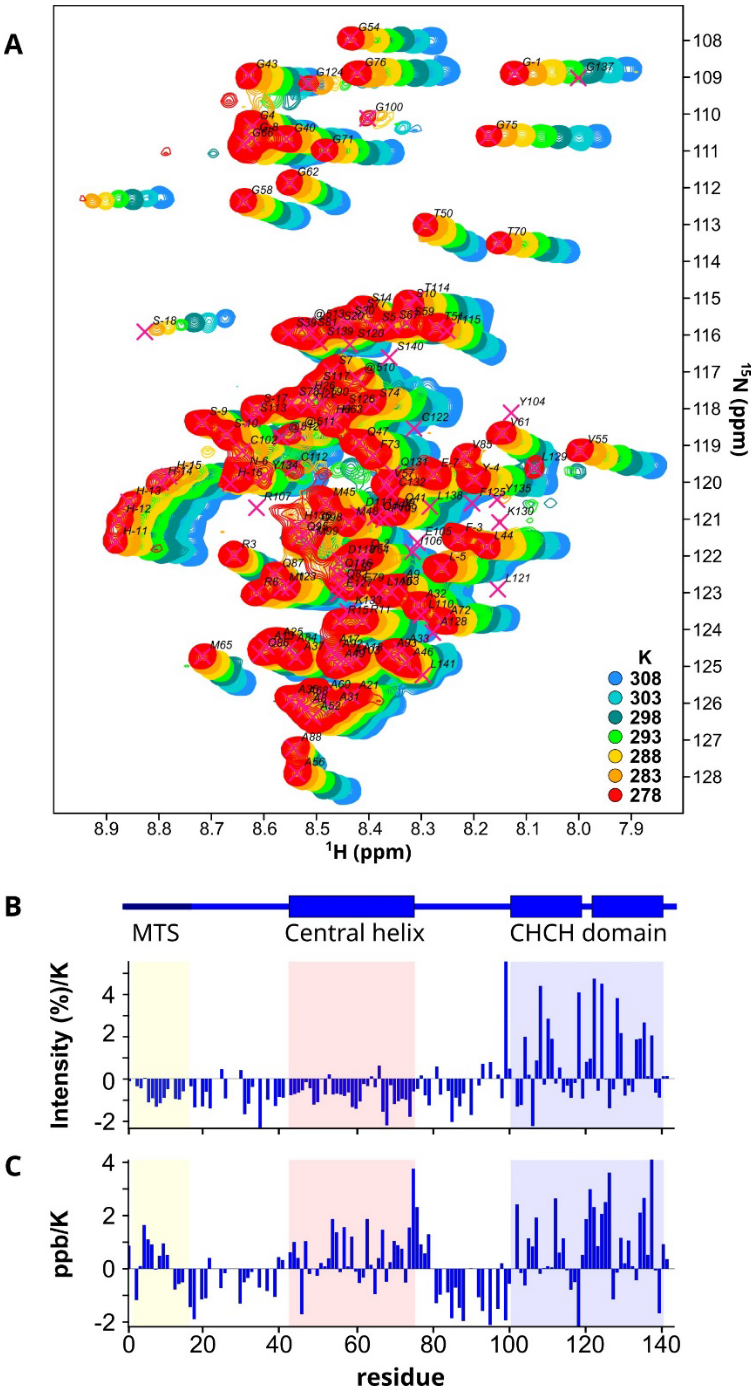


**Supplementary Fig. S6. Temperature-dependent NMR analysis of D10 dynamics. A**, Overlay of $^{1}H$–$^{15}N$ HSQC spectra of full-length D10 recorded over a temperature ramp from 278 to 308 K. Spectra were acquired using 50 µM $^{15}N$-labelled D10 in 20 mM MES pH 6.1, 150 mM NaCl, 0.05% $NaN_3$ and 10% $D_2O$. Progressive changes in peak position and intensity reveal temperature-dependent modulation of D10 dynamics. **B**, Residue-resolved temperature dependence of peak intensity, expressed as intensity change per Kelvin. Several CHCH-domain resonances recover intensity upon heating, consistent with reduced intermolecular exchange and/or weakening of self-association at higher temperature. In contrast, many N-terminal resonances show modest intensity loss, consistent with increased conformational dynamics in the intrinsically disordered region. **C**, Residue-resolved amide proton temperature coefficients. The CHCH domain and adjacent central region show the strongest temperature-sensitive behavior , supporting their contribution to D10 conformational dynamics and reversible self-association.

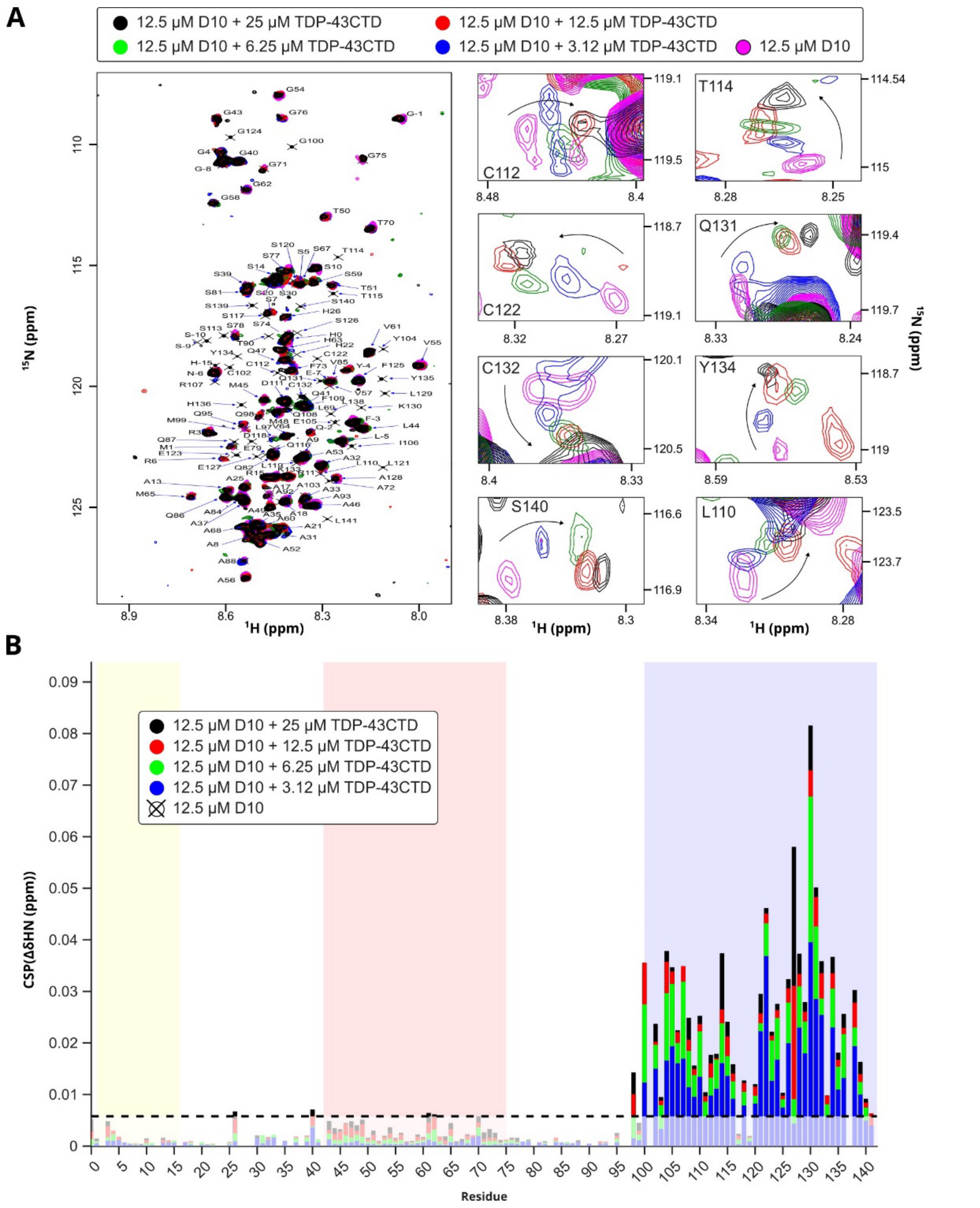


**Supplementary Fig. S7. D10-observed NMR titration maps TDP-43CTD binding to the CHCH domain. A**, Overlay of $^{1}H$–$^{15}N$ HSQC spectra of $^{15}N$-labelled D10 recorded in the absence and presence of increasing concentrations of unlabelled TDP-43CTD. Spectra were acquired using 12.5 µM D10 and 0, 3, 6, 12.5 or 25 µM TDP-43CTD in 20 mM MES pH 6.1, 150 mM NaCl, 0.05% $NaN_3$ and 10% $D_2O$ at 308 K. Representative zooms highlight continuous peak trajectories for CHCH-domain residues L110, T114, C122 and C132, consistent with fast exchange on the NMR timescale. **B**, CSPs in D10 upon addition of TDP-43CTD. Perturbations are strongly enriched within the C-terminal CHCH domain, whereas the N-terminal region and central helical segment remain only weakly affected. The dashed line indicates the threshold used to highlight perturbed residues. Shaded regions indicate the mitochondrial targeting sequence, central helical region and CHCH domain.

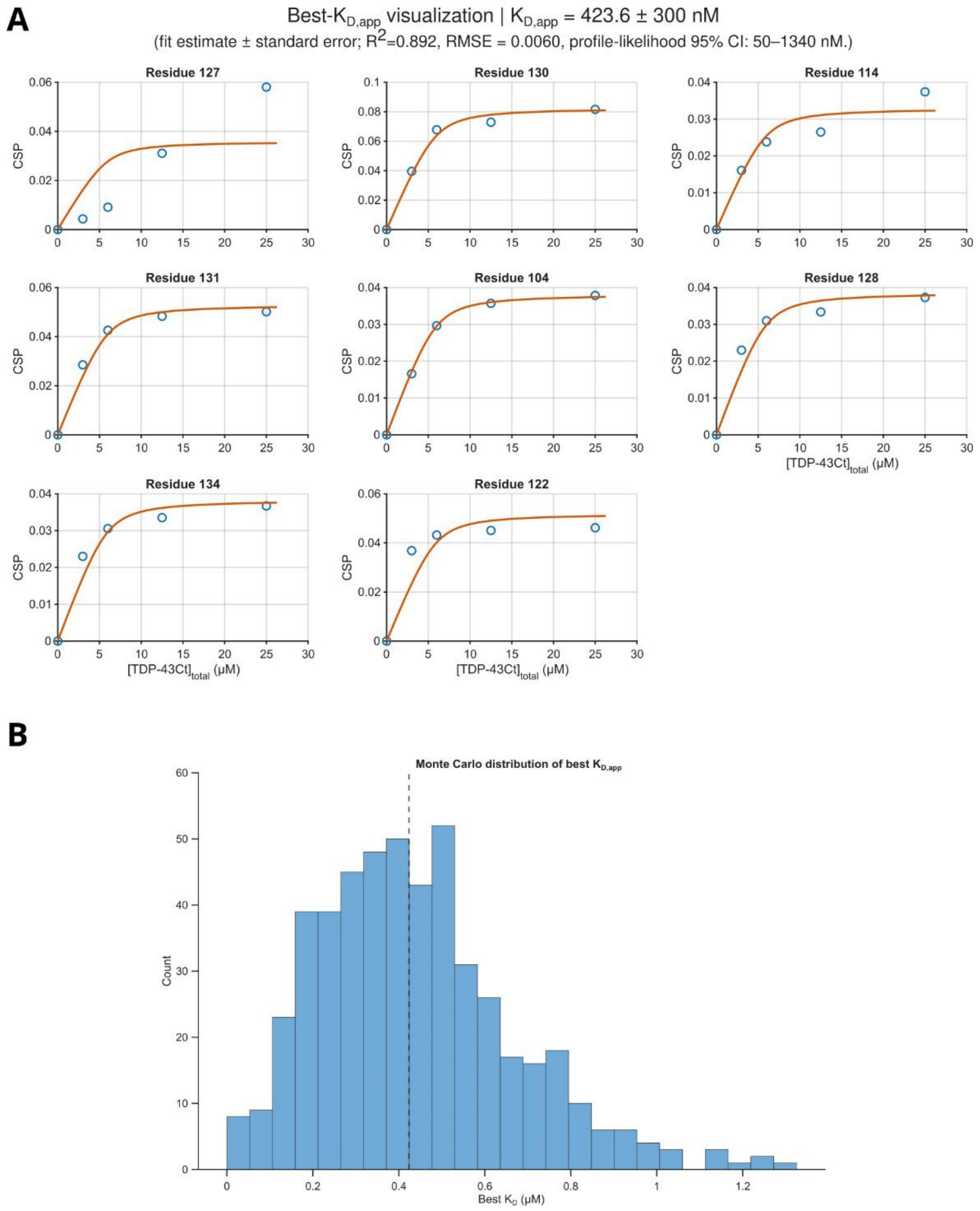


**Supplementary Fig. S8. Apparent affinity estimation from D10-observed NMR titrations. A**, Concentration-dependent CSP profiles for selected D10 residues most responsive to TDP-43CTD addition. Experimental CSP values are shown together with fits to a one-site binding model. The analyzed residues, including R127, K130, T114, Q131, Y104, A128, Y134 and C122, are located within or close to the CHCH-domain interaction surface. The best-fit apparent dissociation constant was $K_D$ = 424 nM. **B**, Monte Carlo distribution of best-fit $K_D$ values obtained from resampling of the D10-observed titration data. The dashed line indicates the best-fit $K_D$. The distribution supports a submicromolar apparent interaction, while reflecting the uncertainty associated with modest CSP amplitudes and a dynamic fast-exchanging complex.

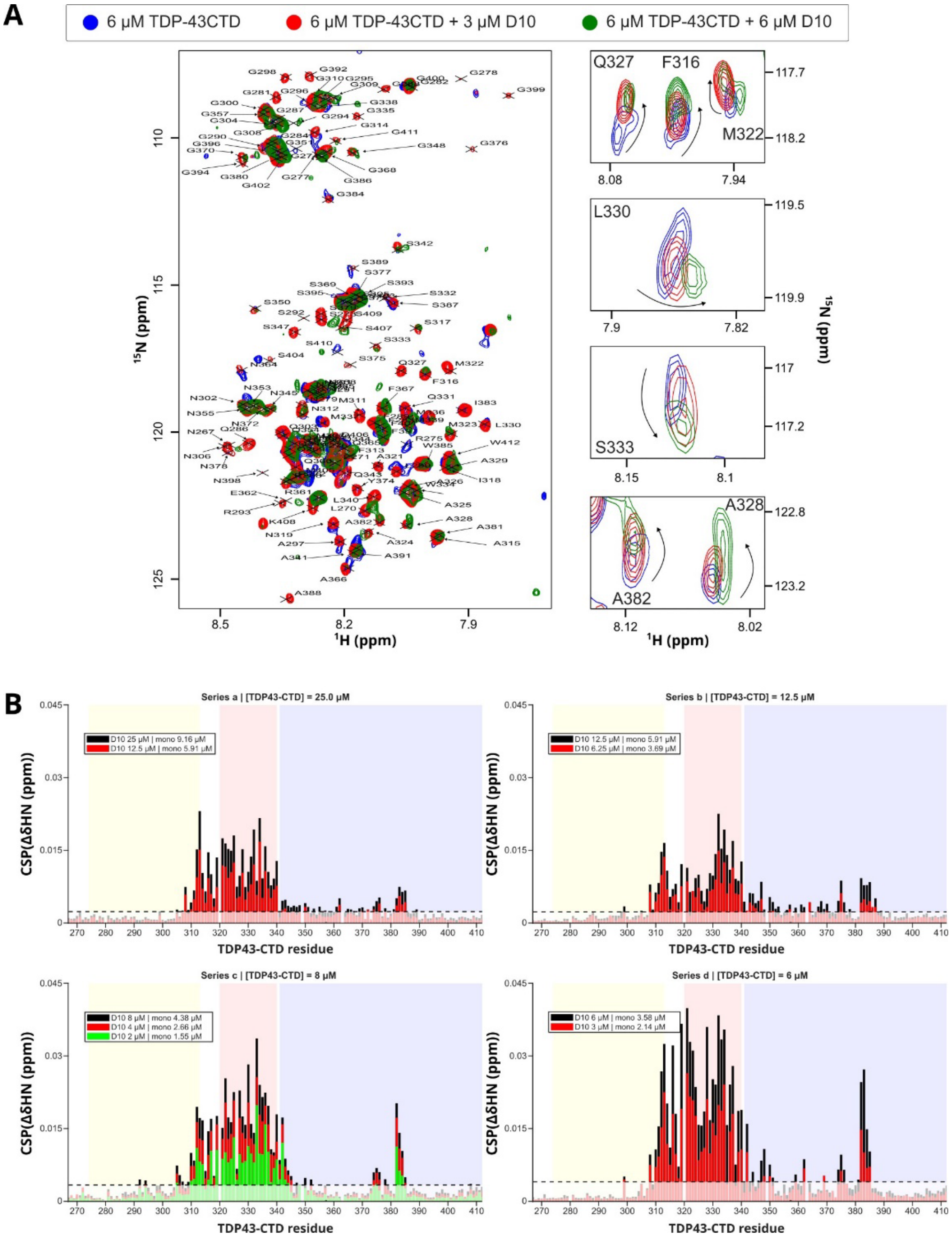


**Supplementary Fig. S9. TDP-43CTD-observed NMR titrations identify the conserved hydrophobic helix as the D10-binding region.** **A**, Overlay of $^{1}H$–$^{15}N$ HSQC spectra of $^{15}N$-labelled TDP-43CTD recorded upon addition of unlabelled D10. Spectra were acquired in 20 mM MES pH 6.1, 150 mM NaCl, 0.05% $NaN_3$ and 10% $D_2O$ at 308 K. Representative zooms show D10-dependent peak movements for residues within the conserved hydrophobic helical region, including F316, M322, Q327, A328, L330 and S333, as well as a distal C-terminal region around A382. Arrows indicate the direction of peak displacement upon D10 addition. **B**, Residue-resolved CSP profiles from independent titration series recorded at different TDP-43CTD concentrations. D10 concentrations were corrected to effective monomeric D10 values using the D10 monomer–dimer equilibrium determined in Fig. 1. Across the different series, CSPs reproducibly cluster within the conserved hydrophobic helical region of TDP-43CTD and its immediate flanking residues, with weaker perturbations detected around the distal C-terminal region.

**A**

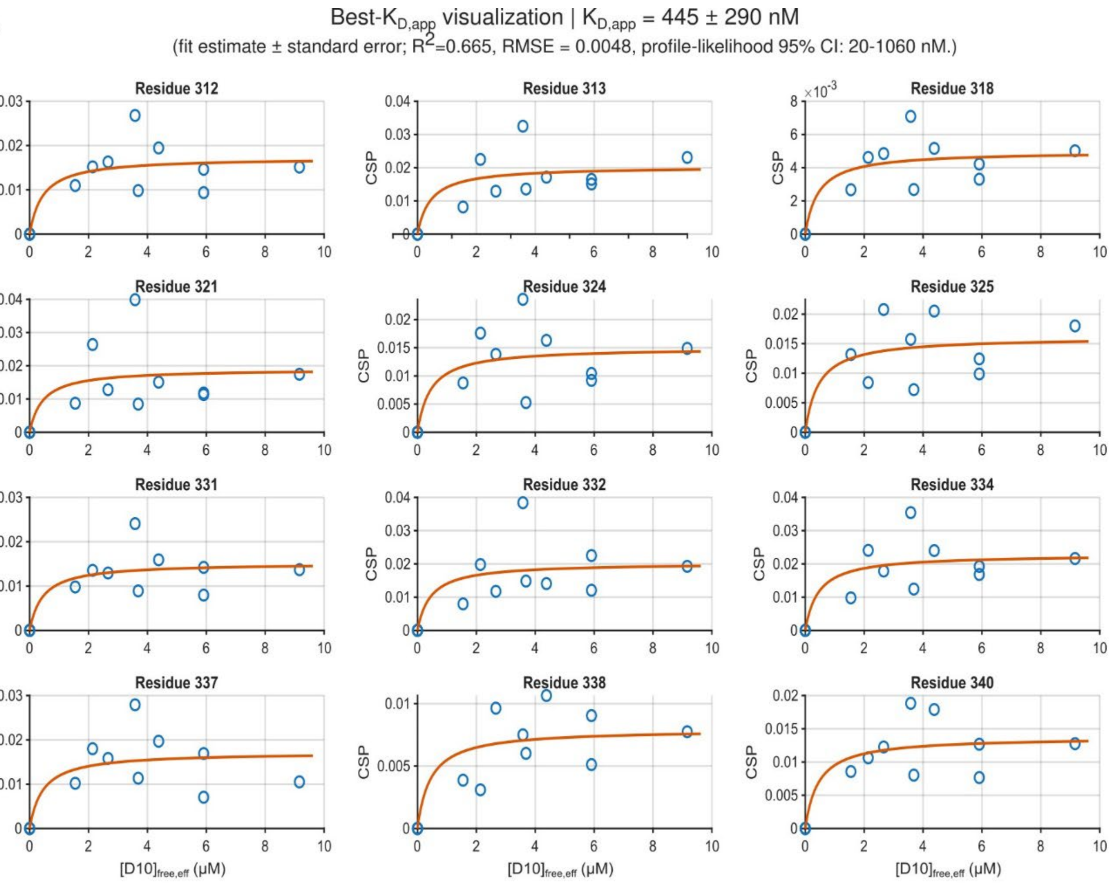


**B**

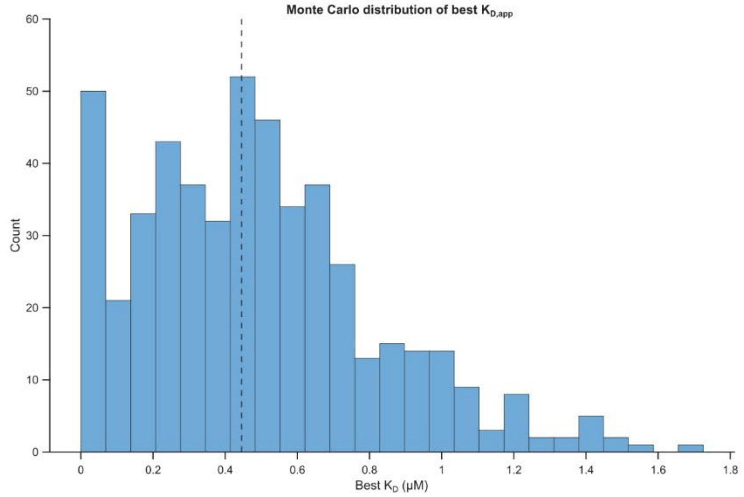


**Supplementary Fig. S10. Apparent affinity estimation from TDP-43CTD-observed NMR titrations. A**, Concentration-dependent CSP profiles for selected TDP-43CTD residues within the D10-responsive region. Experimental CSP values are shown together with fits to a one-site binding model using the estimated effective monomeric D10 concentration. The fitted residues include positions within the conserved hydrophobic helical region, such as residues 312, 313, 318, 321, 324, 325, 331, 332, 334, 337, 338 and 340. The best-fit apparent dissociation constant was $K_D$ = 445 nM. **B**, Monte Carlo distribution of best-fit $K_D$ values obtained from resampling of the TDP-43CTD-observed titration data. The dashed line indicates the best-fit $K_D$. The distribution supports an apparent submicromolar interaction and is consistent with the affinity obtained from the reciprocal D10-observed titration.

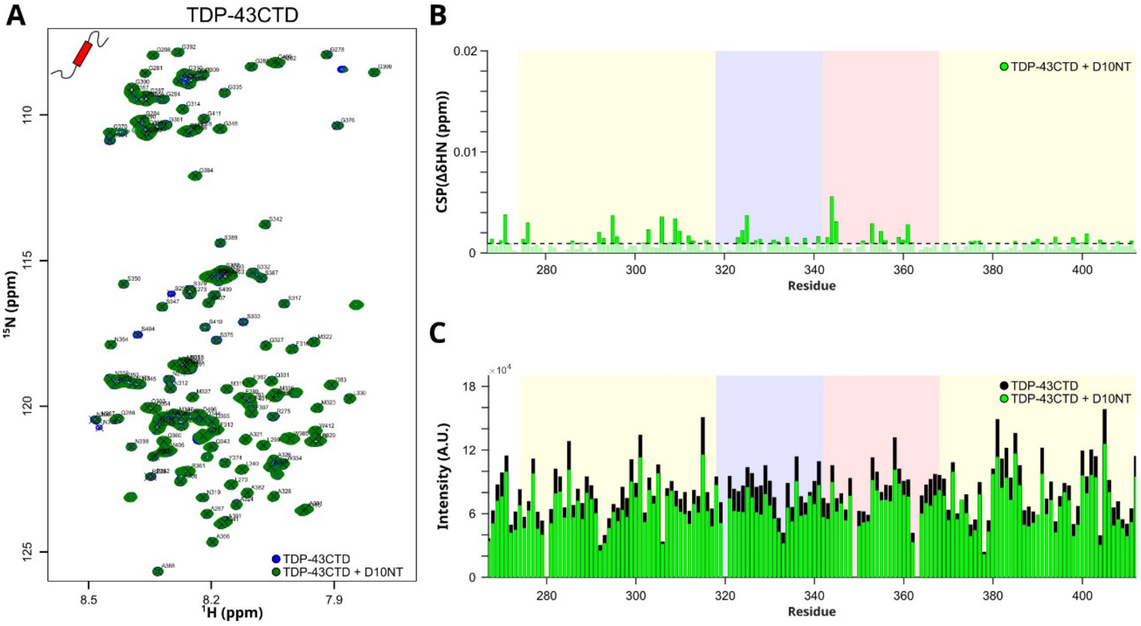


**Supplementary Fig. S11. The D10NT does not reproduce the interaction with TDP-43CTD. A**, Overlay of $^{1}$H–$^{15}$N HSQC spectra of 12.5 µM $^{15}$N-labelled TDP-43CTD recorded alone and after addition of an equimolar amount of D10NT. Spectra were acquired in 20 mM MES pH 6.1, 150 mM NaCl, 0.05% $NaN_3$ and 10% $D_2O$ at 308 K. **B**, CSPs induced by D10NT addition across the TDP-43CTD sequence. Only negligible perturbations are observed, including within the conserved hydrophobic helical region that is strongly perturbed by full-length D10. **C**, Residue-resolved peak intensities of TDP-43CTD in the absence and presence of D10NT. Peak intensities remain broadly preserved across the sequence. Together, these data indicate that the D10NT does not detectably engage TDP-43CTD under these conditions, supporting the conclusion that TDP-43CTD recognition requires the CHCH-containing region of D10.

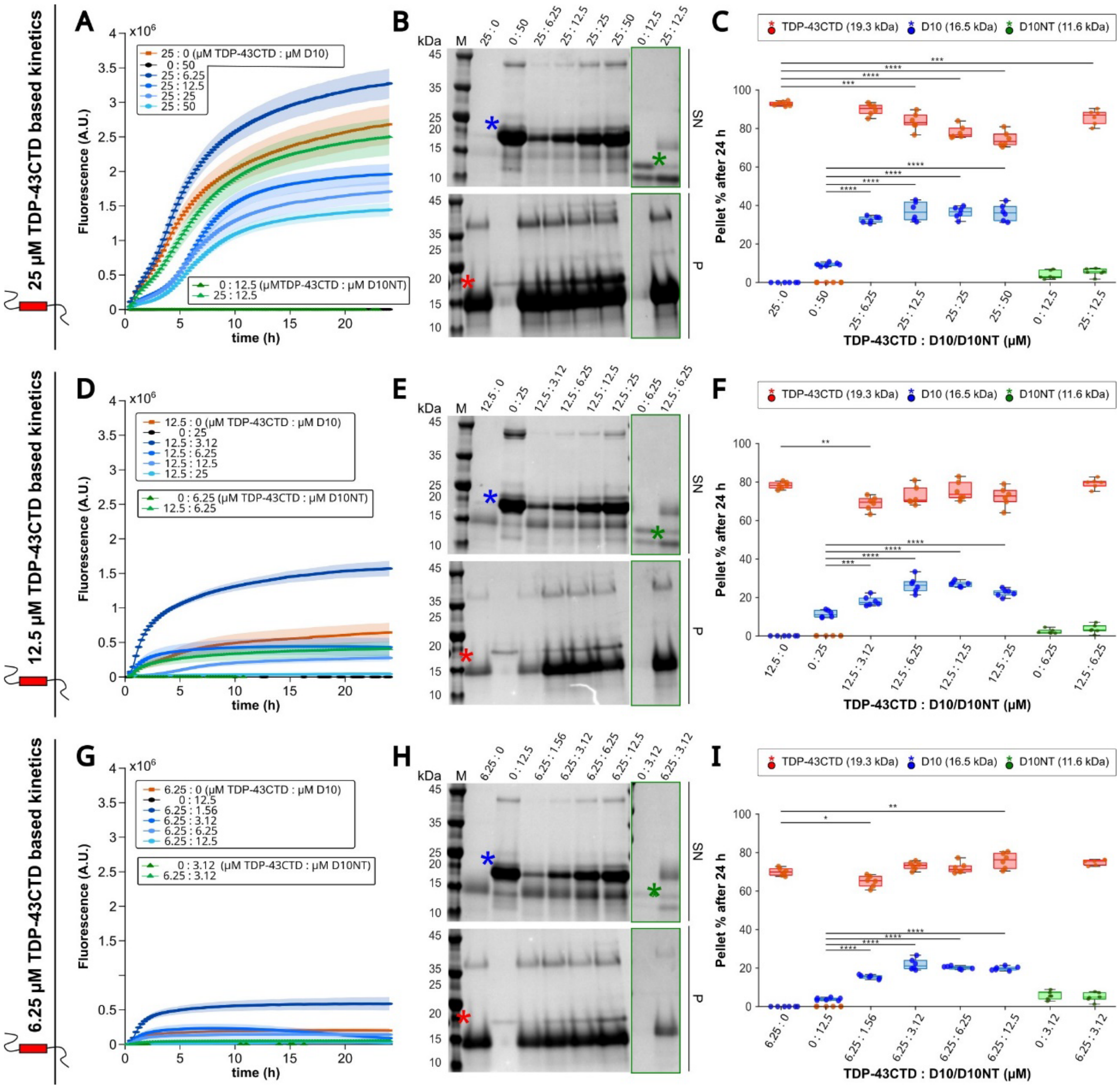


**Supplementary Fig. S12. Concentration-dependent modulation of TDP-43CTD aggregation by D10. A, D, G**, Thioflavin-T aggregation kinetics of TDP-43CTD recorded at 25 µM (A), 12.5 µM (D) or 6.25 µM (G) in the absence or presence of increasing amounts of full-length D10 or the isolated N-terminal construct D10NT. Reactions were prepared at the indicated TDP-43CTD:D10 or TDP-43CTD:D10NT molar ratios in 20 mM MES pH 6.1, 120 mM NaCl, 0.05% $NaN_3$ and 15 µM ThT, and incubated for 24 h at 35 °C under quiescent conditions. Fluorescence was recorded every 15 min. Lines represent mean fluorescence traces and shaded regions indicate variability between replicate measurements. Full-length D10 altered TDP-43CTD aggregation kinetics in a concentration- and stoichiometry-dependent manner, whereas D10NT showed little or no effect. **B, E, H**, Representative SDS-PAGE analysis of non-sedimented/supernatant (SN) and pellet (P) fractions after 24 h of aggregation for reactions containing 25 µM (B), 12.5 µM (E) or 6.25 µM (H) TDP-43CTD. Samples were centrifuged at 16,000 × g for 30 min at 4 °C, and equivalent SN and P fractions were resolved on 4–12% polyacrylamide gels. The red, blue and green asterisks indicate the bands quantified for TDP-43CTD, D10 and D10NT, respectively. The TDP-43CTD band was assigned from purified protein controls and from its absence in D10- or D10NT-only lanes, despite its anomalous apparent electrophoretic mobility. **C, F, I**, Densitometric quantification of the percentage of each protein recovered in the pellet fraction after 24 h for reactions containing 25 µM (C), 12.5 µM (F) or 6.25 µM (I) TDP-43CTD. TDP-43CTD was predominantly recovered in the pellet fraction when incubated alone, whereas D10 showed increased pellet association in the presence of TDP-43CTD. The magnitude of TDP-43CTD redistribution was concentration-dependent and non-linear, consistent with a balance between TDP-43CTD self-assembly, D10 self-association and heterotypic D10–TDP-43CTD interactions. D10NT remained largely soluble and did not reproduce the D10-dependent redistribution. Each point represents an independent replicate; boxes indicate the interquartile range and median. Statistical comparisons were performed relative to the corresponding single-protein or TDP-43CTD-alone controls; $*p < 0.05$, $**p < 0.01$, $***p < 0.001$, $****p < 0.0001$.

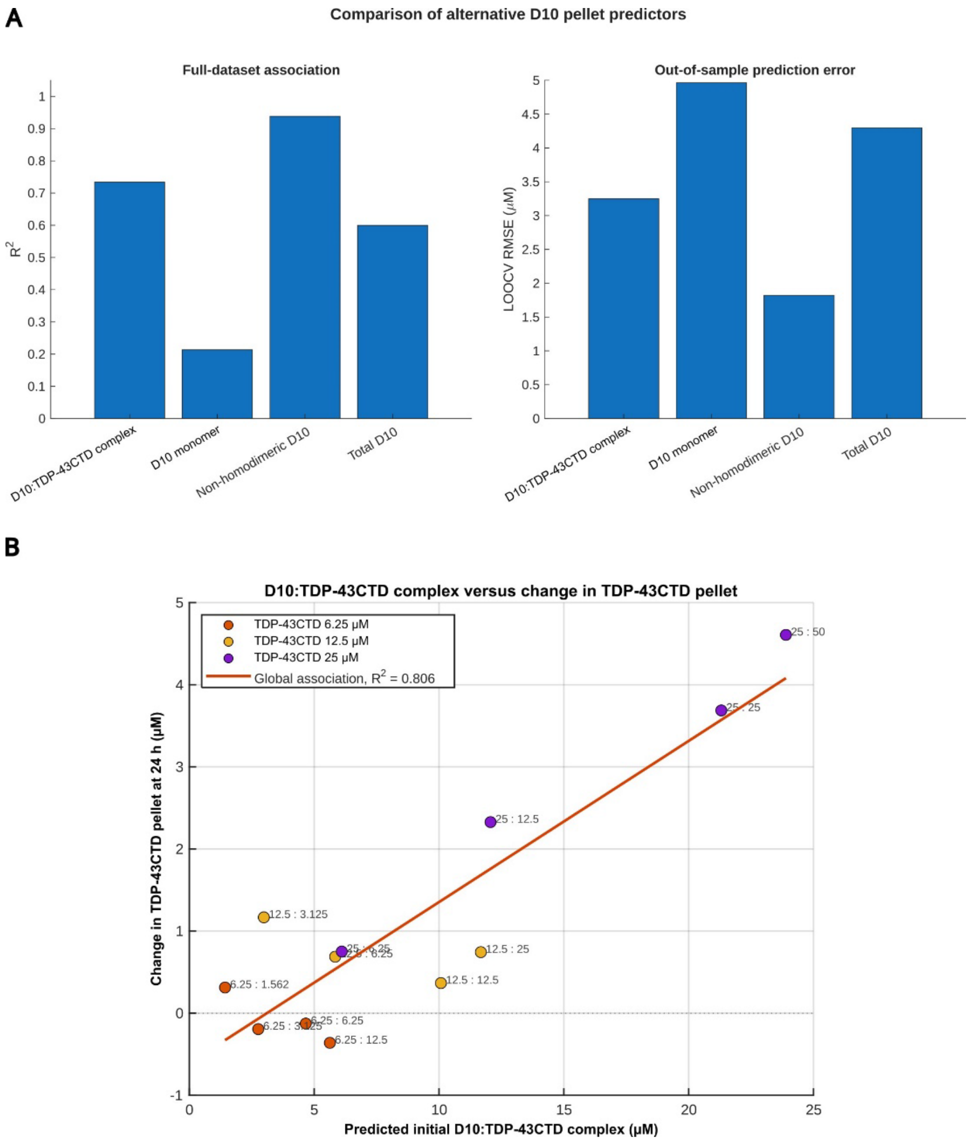


**Supplementary Figure S13. Equilibrium populations calculated from the explicit cubic-equation formulation identify the non-homodimeric D10 pool as the strongest predictor of D10 sedimentation. A,** Comparison of alternative predictors of D10 recovery in the pellet after 24 h. Initial soluble populations were calculated for each experimental condition using the explicit cubic-equation formulation describing D10 homodimerization and formation of a 1:1 D10:TDP-43CTD complex. The model used the apparent dissociation constants $K_{D,D10}$=10.6 μM and $K_{D,DT}$=0.434 μM, with TDP-43CTD treated as predominantly monomeric at the beginning of the reaction. Left, full-dataset explanatory performance, expressed as the coefficient of determination ($R^2$), for linear associations between each predictor and the experimentally measured D10 pellet concentration. Right, out-of-sample prediction error obtained by leave-one-condition-out cross-validation and expressed as RMSE. The tested predictors were the cubic-equation-derived D10:TDP-43CTD complex, free D10 monomer and non-homodimeric D10 pool ($[D10]_{monomer}$+[D10:TDP-43CTD]), together with total D10 concentration. The non-homodimeric D10 pool showed the strongest full-dataset association ($R^2$=0.938) and the lowest cross-validated prediction error (*RMSE* = 1.82 μM). **B**, Association between the initial D10:TDP-43CTD complex population calculated from the explicit cubic equation and the change in TDP-43CTD recovered in the pellet after 24 h. The change in pellet recovery was calculated relative to the matched TDP-43CTD-only control as $\Delta TDP\text{-}43CTD_{pellet}=TDP\text{-}43CTD_{pellet,control}-TDP\text{-}43CTD_{pellet,+D10}$. Positive values indicate reduced TDP-43CTD pellet recovery in the presence of D10, whereas negative values indicate increased recovery. Points are colored according to total TDP-43CTD concentration, and labels indicate the corresponding TDP-43CTD:D10 concentrations in μM. The red line represents the global linear association across all conditions ($R^2$=0.806). Because this relationship was strongest at 25 μM TDP-43CTD and weaker and non-monotonic at lower concentrations, it was interpreted as a concentration-dependent endpoint association rather than as a general kinetic model of TDP-43CTD aggregation. The calculated populations describe the initial soluble equilibrium and do not reconstruct the pathway leading to the 24-h sedimentation endpoint.

**Independent numerical validation using a damped-Newton solution of the coupled D10 - TDP-43CTD mass-balance equations**

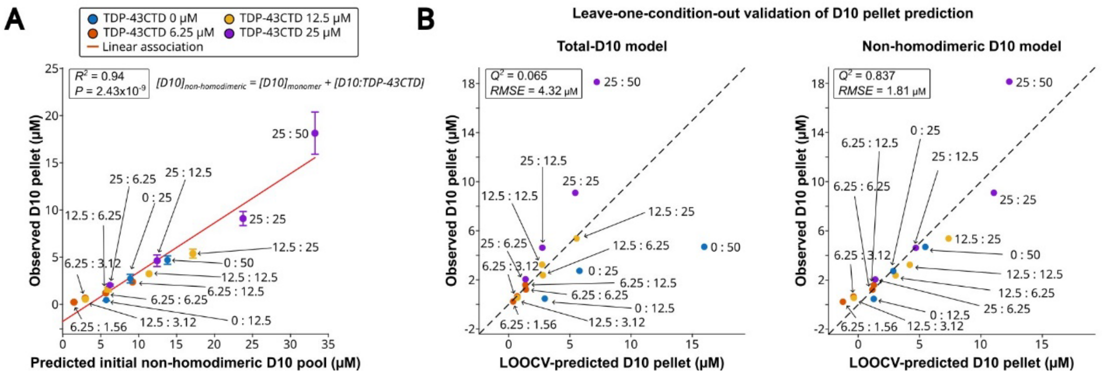


**Supplementary Figure S14. Independent numerical validation of the relationship between the initial non-homodimeric D10 population and D10 sedimentation.** Initial equilibrium populations were calculated by numerically solving the complete coupled D10–TDP-43CTD mass-balance equations using a damped-Newton algorithm. The model included D10 monomer–homodimer equilibrium, formation of the 1:1 D10:TDP-43CTD complex and the reported TDP-43CTD monomer–dimer–tetramer self-association pathway. The equations were solved simultaneously for the free D10 and TDP-43CTD concentrations in each experimental condition. **A**, Association between the numerically calculated initial non-homodimeric D10 population and the experimentally observed amount of D10 recovered in the pellet after 24 h. The non-homodimeric population was defined as the sum of free D10 monomer and heterotypically bound D10, $[D10]_{non\text{-}homodimeric}=[D10]_{monomer}+[D10:TDP\text{-}43CTD]$. Each point represents one initial TDP-43CTD:D10 condition and is coloured according to total TDP-43CTD concentration; labels indicate total TDP-43CTD:D10 concentrations in µM. Error bars show variability in the experimentally measured D10 pellet concentration. The red line represents the linear association over the experimentally sampled range ($R^2$=0.940, $P$=2.43×10$^{-9}$). **B**, Leave-one-condition-out cross-validation of linear models predicting D10 pellet recovery from either total D10 concentration or the numerically calculated initial non-homodimeric D10 population. For each point, the indicated condition was excluded, the model was fitted to all remaining conditions and the pellet concentration of the excluded condition was predicted. Dashed lines indicate identity between predicted and observed values. The non-homodimeric D10 model showed substantially greater out-of-sample performance ($Q^2$=0.837, *RMSE* = 1.81 µM) than the total-D10 model ($Q^2$=0.065, *RMSE* = 4.32 µM). The numerical solution reproduced the relationship obtained using the explicit cubic-equation formulation in the main analysis, supporting the conclusion that the minor initial TDP-43CTD dimeric and tetrameric populations do not materially affect the calculated D10-state distribution or its association with the 24-h sedimentation endpoint. These calculations describe the initial soluble equilibrium populations and do not constitute a kinetic reconstruction of pellet formation.

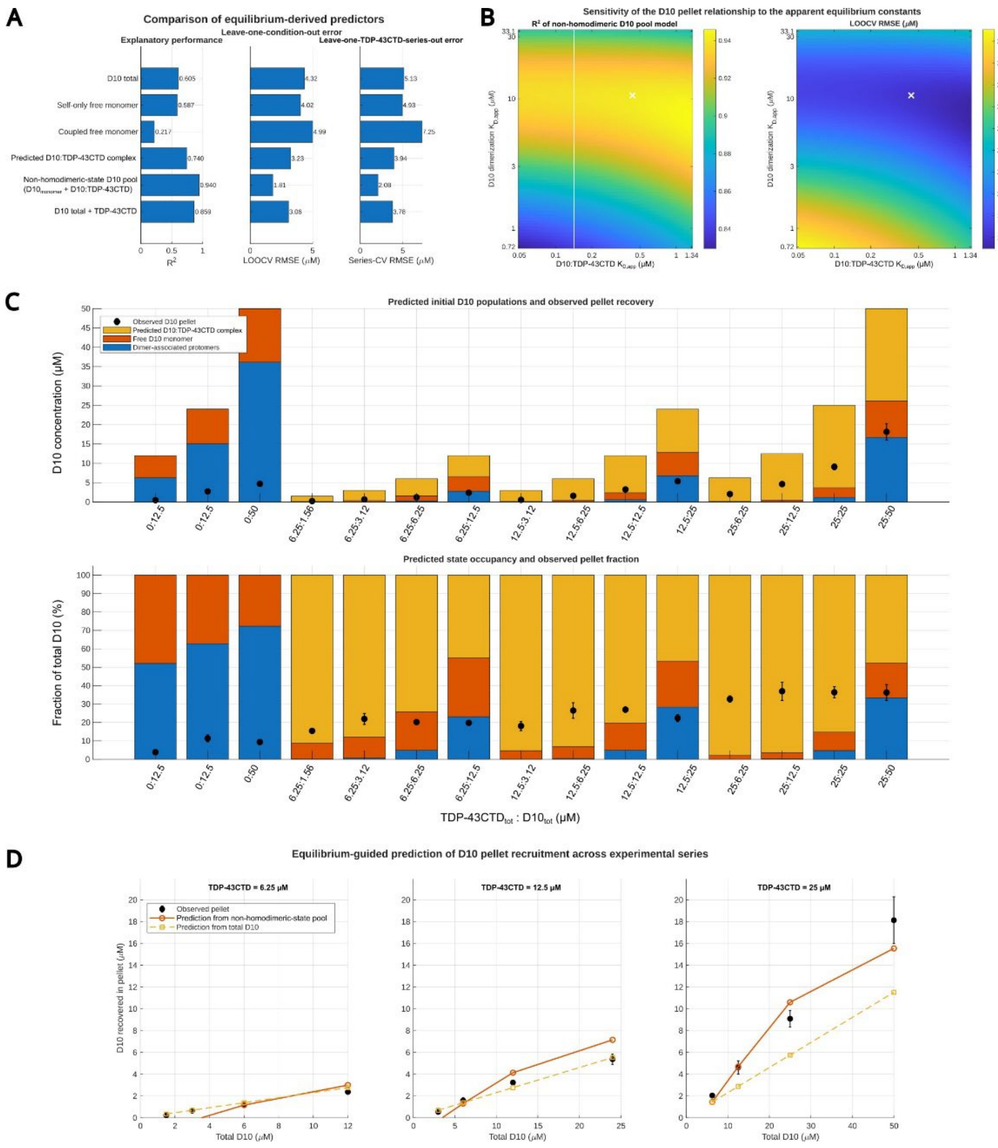


**Supplementary Fig. S15. Equilibrium-derived predictors of D10 pellet recruitment and their robustness to uncertainty in the apparent equilibrium constants. A,** Comparison of alternative equilibrium-derived predictors of D10 recovery in the pellet fraction after 24 h. Bars show the explanatory performance of each predictor in the full dataset ($R^2$), together with prediction errors obtained by leave-one-condition-out cross-validation (LOOCV RMSE, µM) and by leave-one-TDP-43CTD-series-out validation (Series-CV RMSE, µM). The tested predictors were total D10 concentration, free D10 monomer predicted from the D10 self-association equilibrium alone ("self-only free monomer"), free D10 monomer predicted in the coupled homo-/heterotypic system ("coupled free monomer"), the predicted D10–TDP-43CTD complex, the combined non-homodimeric D10 pool ($[D10]_{monomer}$ + [D10–TDP-43CTD]) and the sum of total D10 plus total TDP-43CTD concentrations. The non-homodimeric D10 pool provided the strongest explanatory and predictive performance. **B,** Sensitivity of the non-homodimeric D10 pool relationship to the apparent equilibrium constants used in the population calculation. Heat maps show the resulting $R^2$ values (left) and LOOCV RMSE values (right) across the tested ranges of the D10 homodimerization constant and the D10–TDP-43CTD binding constant. The white cross marks the central fitted values used in the main analysis ($K_{D,app}$ = 10.6 µM for D10 self-association and $K_{D,app}$ ≈ 0.43 µM for the D10–TDP-43CTD interaction). The relationship remained robust across the uncertainty ranges of both parameters. **C,** Predicted initial D10 state populations and observed pellet recovery for each experimental condition. Top, stacked bars show the calculated concentrations of dimer-associated D10 protomers (blue), free D10 monomer (yellow) and the predicted D10–TDP-43CTD complex (orange), together with the experimentally observed amount of D10 recovered in the pellet after 24 h (black symbols, mean ± variability). Bottom, the same data represented as fractions of total D10. Conditions are indicated as total TDP-43CTD : total D10 concentrations (µM). These plots illustrate the redistribution of D10 from homodimer-associated states towards free and heterotypically bound states as TDP-43CTD concentration increases. **D,** Equilibrium-guided prediction of D10 pellet recruitment across the three experimental TDP-43CTD series (6.25, 12.5 and 25 µM TDP-43CTD). Black symbols show the observed D10 recovered in the pellet after 24 h. Orange lines show predictions from the non-homodimeric D10 pool model, whereas dashed yellow lines show predictions from the model based on total D10 concentration alone. The non-homodimeric-state model more closely reproduced the observed concentration dependence across the experimental series.

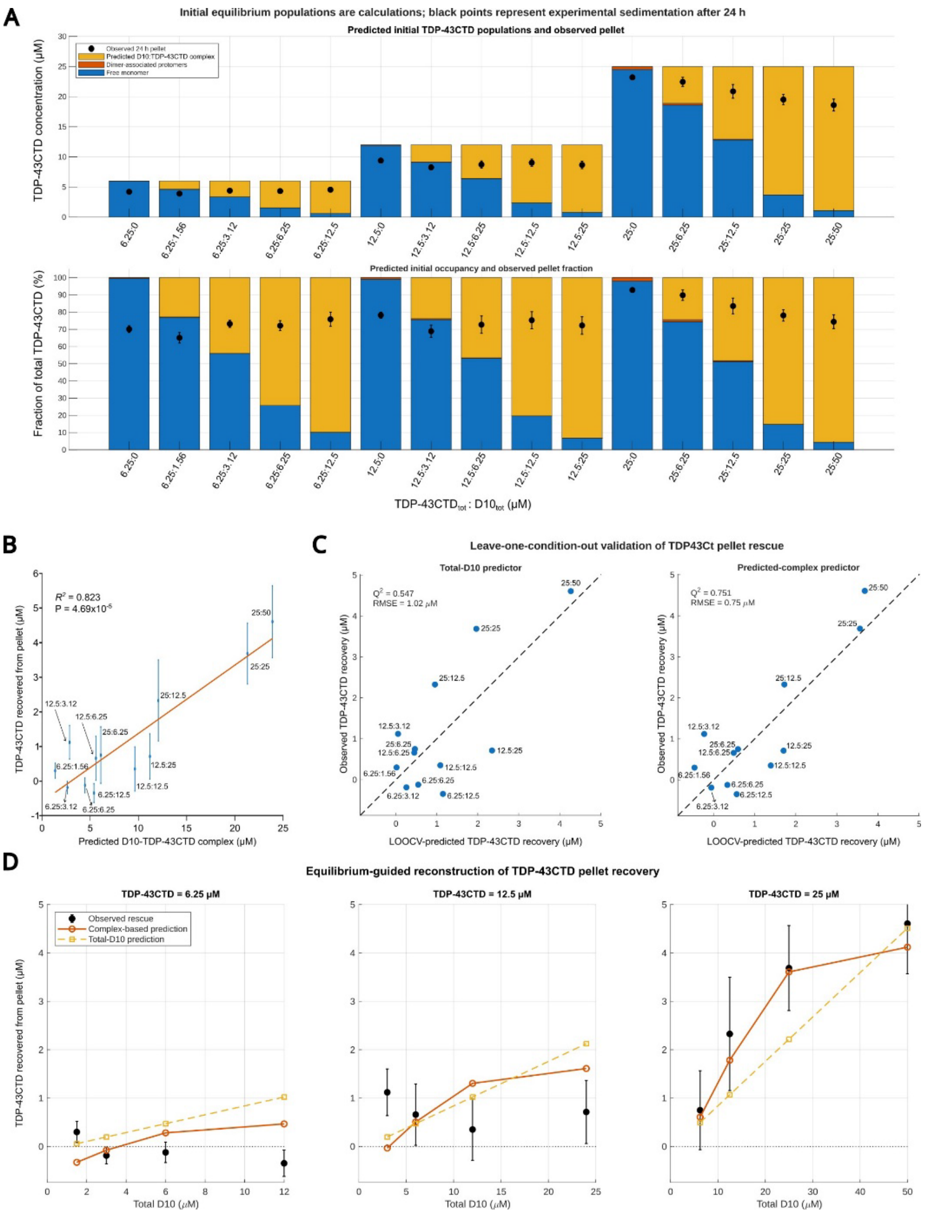


**Supplementary Fig. S16. Equilibrium-derived D10–TDP-43CTD complex populations predict partial rescue of TDP-43CTD from the pellet. A,** Predicted initial TDP-43CTD populations and experimentally observed pellet recovery after 24 h for each TDP-43CTD:D10 condition. Top, stacked bars show the calculated concentrations of free TDP-43CTD monomer, dimer-associated TDP-43CTD protomers and heterotypic D10–TDP-43CTD complex. Black symbols indicate the experimentally measured amount of TDP-43CTD recovered in the pellet after 24 h. Bottom, the same predicted populations and experimental pellet values expressed as fractions of total TDP-43CTD. Conditions are labelled as total TDP-43CTD:total D10 concentrations in µM. The calculations predict that TDP-43CTD is predominantly monomeric in the absence of D10, with only a minor homodimer-associated population, whereas increasing D10 progressively redistributes TDP-43CTD into the heterotypic complex. **B,** Relationship between the predicted initial D10–TDP-43CTD complex population and the amount of TDP-43CTD rescued from the pellet, defined as the reduction in pellet-associated TDP-43CTD relative to the corresponding D10-free control. Each point represents one experimental condition, labelled by the initial TDP-43CTD:D10 concentrations in µM. The orange line represents the linear regression ($R^2$ = 0.823, $P = 4.69 \times 10^{-5}$), and error bars indicate variability in the experimentally determined rescue values. **C,** Leave-one-condition-out cross-validation of models predicting TDP-43CTD rescue from either total D10 concentration or the calculated initial D10–TDP-43CTD complex population. Dashed lines indicate perfect agreement between predicted and observed values. The predicted-complex model showed greater predictive performance ($Q^2$ = 0.751; RMSE = 0.75 µM) than the total-D10 model ($Q^2$ = 0.547; RMSE = 1.02 µM). **D,** Equilibrium-guided reconstruction of TDP-43CTD rescue across the experimental series containing 6.25, 12.5 or 25 µM TDP-43CTD. Black symbols show the experimentally observed rescue from the pellet, solid orange lines show predictions from the D10–TDP-43CTD complex model and dashed yellow lines show predictions based on total D10 concentration alone. The complex-based model more closely reproduced the observed response, particularly at 25 µM TDP-43CTD, whereas the weaker and non-monotonic effects observed at lower TDP-43CTD concentrations were less completely captured by either predictor.

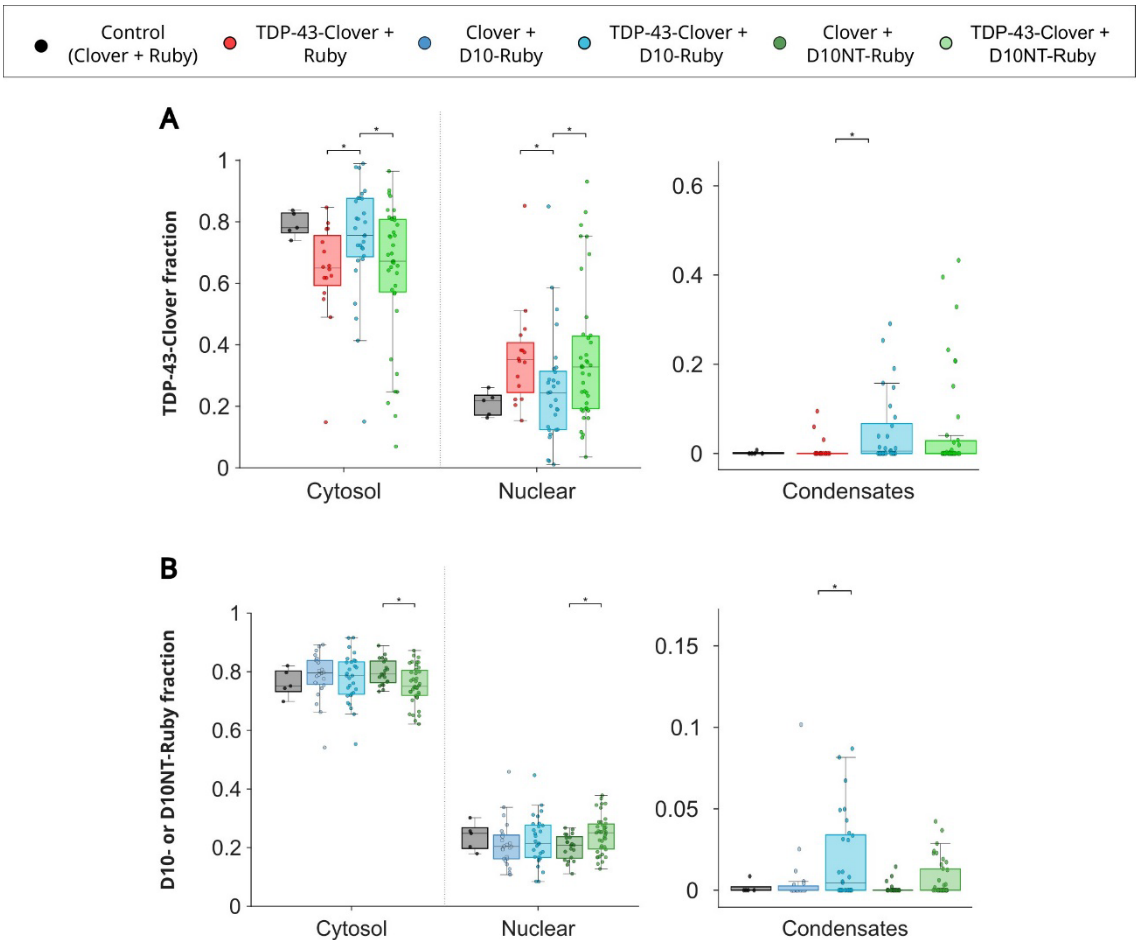


**Supplementary Fig. S17. Compartmental distribution of Clover- and Ruby-tagged proteins in HeLa cells. A**, Quantification of the compartmental distribution of the Clover signal across cytosolic, nuclear and condensate-associated regions in HeLa cells transiently expressing the indicated constructs. Conditions included Clover/Ruby control, TDP-43-Clover/Ruby, Clover/D10-Ruby, TDP-43-Clover/D10-Ruby, Clover/D10NT-Ruby and TDP-43-Clover/D10NT-Ruby. Co-expression of full-length D10-Ruby altered the cytosolic/nuclear distribution of TDP-43-Clover compared with TDP-43-Clover/Ruby, whereas D10NT-Ruby produced a weaker effect. Condensate-associated fractions are shown on a separate y-axis because of their lower overall values. **B**, Quantification of the compartmental distribution of the Ruby signal corresponding to full-length D10-Ruby or D10NT-Ruby across cytosolic, nuclear and condensate-associated regions. TDP-43-Clover co-expression did not markedly alter the bulk cytosolic/nuclear distribution of full-length D10-Ruby, whereas D10NT-Ruby showed a modest shift towards the nuclear compartment upon TDP-43-Clover co-expression. Cells were fixed 18–24 h after transfection and imaged by confocal microscopy. Regions of interest were defined for cytosolic, nuclear and condensate-associated compartments after background subtraction. Boxes indicate the interquartile range and median, and individual points represent cell-level measurements. Statistical comparisons were performed using Wilcoxon tests with FDR correction; *$p < 0.05$, **$p < 0.01$, ***$p < 0.001$, ****$p < 0.0001$.

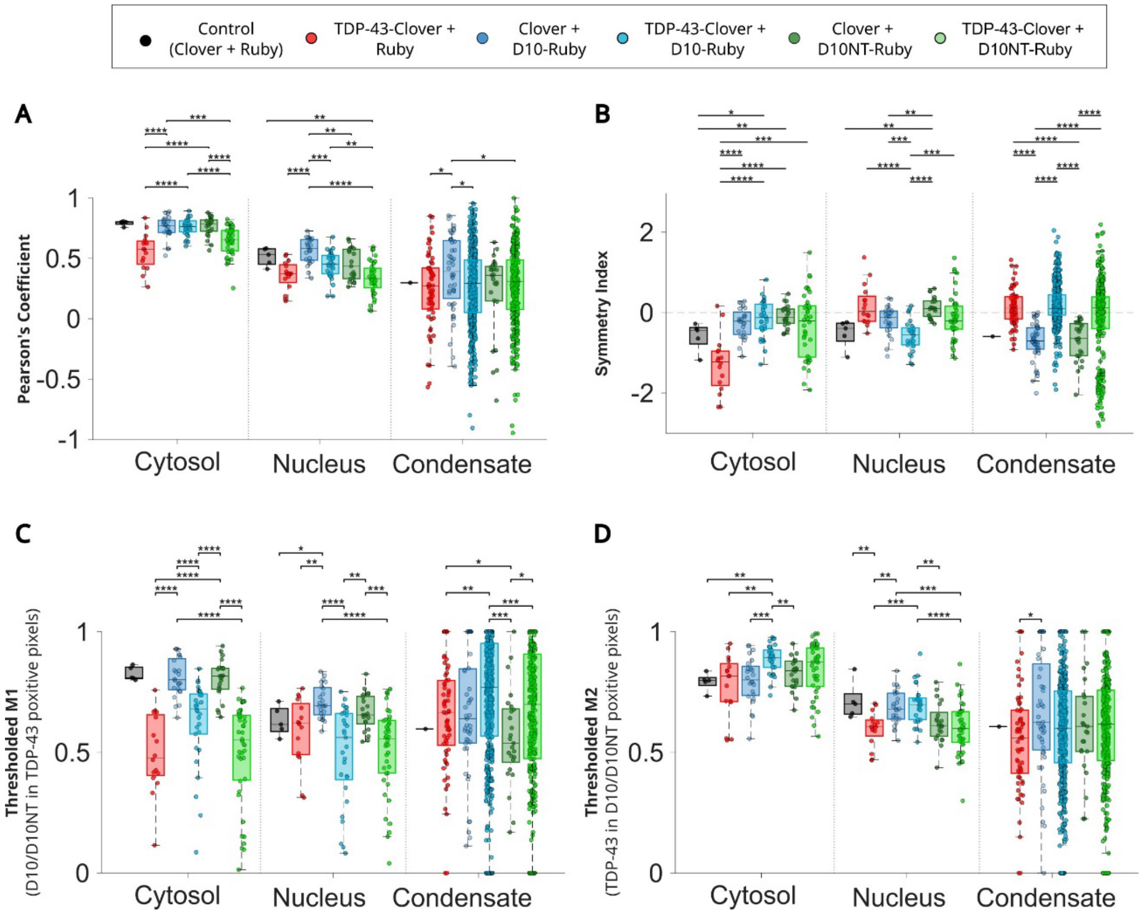


**Supplementary Fig. S18. Extended compartment-resolved colocalization analysis of D10 and TDP-43 in cells. A**, Pearson's correlation coefficients calculated between Clover and Ruby fluorescence signals in cytosolic, nuclear and condensate-associated regions for the indicated transfection conditions. Full-length D10-Ruby showed stronger spatial correlation with TDP-43-Clover than D10NT-Ruby in cytosolic and nuclear compartments, whereas condensate-associated regions showed greater heterogeneity. **B**, Symmetry index derived from directional Manders overlap coefficients, indicating the balance between reciprocal Clover-to-Ruby and Ruby-to-Clover overlap across compartments. Differences in nuclear and condensate-associated regions support a compartment-dependent asymmetry in D10/TDP-43 spatial coupling. **C**, Thresholded Manders M1 analysis, reporting the fraction of D10-Ruby or D10NT-Ruby signal overlapping TDP-43-Clover-positive pixels. Full-length D10 showed increased overlap with TDP-43-positive regions compared with D10NT, particularly in cytosolic and condensate-associated compartments. **D**, Thresholded Manders M2 analysis, reporting the reciprocal fraction of TDP-43-Clover signal overlapping D10-Ruby- or D10NT-Ruby-positive pixels. The strongest CHCH-dependent difference was detected in the nuclear compartment. Colocalization analyses were performed in Fiji/ImageJ using JaCoP after background subtraction. Cytosolic, nuclear and condensate-associated regions of interest were analyzed separately, low-quality ROIs were excluded, and measurements were aggregated at the cell level before statistical testing. Boxes indicate the interquartile range and median, and individual points represent cell-level measurements. Statistical comparisons were performed using Wilcoxon tests with FDR correction; *$p < 0.05$, **$p < 0.01$, ***$p < 0.001$, ****$p < 0.0001$.
